# Noise-referenced multi-shell diffusion-weighted imaging for regional cerebrospinal fluid thermometry

**DOI:** 10.64898/2026.10.28.762634

**Authors:** Christian Behler, Michael Breakspear, Saadallah Ramadan, Thuy Thanh Dao, Thomas G. Close, Sjoerd B. Vos, Renate Thienel, Joseph Giorgio, Anna Behler, Michael Fay

**Author notes:** Joint senior authors (Anna Behler, Michael Fay).

## Abstract

We developed a multi-shell diffusion-weighted imaging approach for separate ventricular and subarachnoid cerebrospinal fluid thermometry. The approach uses measured image noise to assess agreement among temperature estimates. In 12 participants, temperature was estimated from four non-zero diffusion weightings. For each voxel, disagreement between these estimates was compared with that expected from image noise, accounting for their shared reference image. Voxels meeting this consistency criterion were retained for regional analysis. Mean diffusion-derived temperature (± SD) was 38.4 ± 0.4 °C in the lateral ventricles and 36.0 ± 1.1 °C in the subarachnoid space. Ventricular estimates were consistently higher than subarachnoid estimates across all participants, with a mean difference of 2.4 ± 1.2 °C (95% confidence interval 1.6–3.1 °C). This difference persisted under alternative noise models and temperature estimators. Subarachnoid estimates increased with retained voxel count; in a post hoc analysis of the six participants with the largest retained counts, ventricular estimates remained higher by 1.6 ± 0.7 °C. Absolute temperature accuracy and the physiological origin of the regional difference require independent validation. This approach provides an internal consistency check for CSF thermometry and supports separate assessment of ventricular and subarachnoid temperatures.

## 1. Introduction

Brain temperature reflects the balance between metabolic heat production, cerebral blood flow, and heat exchange with the body and environment (Behler et al., 2026). Implanted probes have shown that brain tissue can be warmer than systemic core temperature, although invasive measurements are generally obtained in neurosurgical or neurocritical care settings (Mariak et al., 1999; Wang et al., 2016). In healthy adults, magnetic resonance spectroscopy (MRS) has demonstrated regional variation in brain temperature. Rzechorzek et al. (2022) reported a mean brain-tissue temperature near 38.5 °C and a mean maximal spatial range of approximately 2.4 °C, with the highest temperatures in the thalamus. These findings show that intracranial temperature varies across regions. Measuring this variation non-invasively could help investigate its relationship to cerebral metabolism and heat exchange. Interpreting regional estimates requires distinguishing temperature differences from measurement inconsistency.

Diffusion-weighted imaging (DWI) thermometry uses the temperature dependence of water diffusion (Mills, 1973). Converting diffusivity to absolute temperature using a free-water calibration requires approximately free diffusion. Intracranial applications have therefore focused on cerebrospinal fluid (CSF), where diffusion is approximately free (Kozak et al., 2010; Tofts et al., 2008). Previous DWI thermometry studies have typically estimated ventricular temperature from a single non-zero diffusion weighting. These temperature measurements have shown associations with parenchymal temperature, age, cognition, and clinical state (Asfuroğlu et al., 2022; Horiuchi et al., 2023; Sakai et al., 2011; Shibukawa et al., 2023; Sumida et al., 2016). Subsequent work investigated b-value selection and motion compensation (Shibukawa et al., 2020), and calibrated the relationship between diffusivity and temperature using an artificial-CSF phantom (Shibukawa et al., 2021). Shibukawa et al. (2021) extended DWI thermometry beyond the ventricles but reported ventricular and subarachnoid CSF as a pooled intracranial estimate rather than as separate regional estimates.

Extending DWI thermometry to the subarachnoid space is challenging because this compartment is thin and therefore vulnerable to tissue partial-volume effects. CSF bulk motion can also alter the diffusion signal (Kozak et al., 2010; Shibukawa et al., 2020). Both effects can bias diffusion-derived temperature. In ocular vitreous thermometry, Derakhshan et al. (2020) showed that apparent temperature depended on b-value sampling and was strongly affected by motion. These findings motivate measurements at multiple diffusion weightings to assess the consistency of temperature estimates. Such an assessment must account for the disagreement expected from measurement noise.

We developed a noise-referenced multi-shell approach to assess agreement among CSF temperature estimates and obtain separate ventricular and subarachnoid estimates. For each voxel, we compared the disagreement among four shell-wise temperature estimates with that expected from measured image noise, accounting for their shared b = 0 reference. We applied the approach to anatomically identified lateral ventricular and subarachnoid compartments, compared regional estimates with conventional single-shell thermometry, and examined sensitivity to tissue admixture, shell selection, and sub-arachnoid sampling.

## 2. Theory

### 2.1 Temperature from the apparent diffusion coefficient

All calculations below are performed voxel-wise. The apparent diffusion coefficient (ADC) at shell *i* is estimated from the mean reference signal *S*_0_ and mean shell signal *S*_i_ as

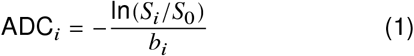

Here, *i* = 1, …, *K* indexes the non-zero diffusion shells (*K* = 4, with *b*_*i*_ ∈ {200, 500, 700, 1100} s/mm^2^). *S*_0_ is averaged over *N*_0_ b = 0 volumes, and *S*_*i*_ over the *N*_*i*_ volumes in shell *i*. For freely diffusing water, the Arrhenius relationship converts ADC to a shell-wise temperature estimate (Mills, 1973):

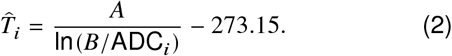

The calibration-specific constants *A* and *B* have units of kelvin and mm^2^/s, respectively. We used the published artificial cerebrospinal fluid (ACSF) calibration (Shibukawa et al., 2021) specified in Section 3.4; sensitivity to the Mills water calibration is reported in Supplementary Section S1. The primary voxel-wise temperature estimate is the median across shells:

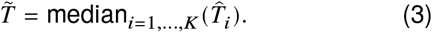

### 2.2 Noise propagation

Marchenko–Pastur principal component analysis (MP-PCA) denoising (Veraart et al., 2016) provides a voxel-wise estimate of the input magnitude-noise standard deviation. Let *σ*_ref_ denote this reference after the same spatial intensity scaling and regridding as the signal, and *a*_*j*_ the actual multiplicative drift factor applied to volume *j*. We use a first-order Gaussian approximation appropriate at high signal-to-noise ratio (Gudbjartsson & Patz, 1995), with zero-centred, independent input-volume errors. For a mean over a set *J* of *N* volumes, its reference noise standard deviation is

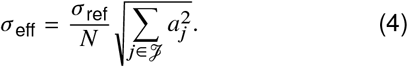

This reduces to 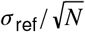 when all *a*_*j*_ = 1. Using the b = 0 and shell-specific volume sets gives

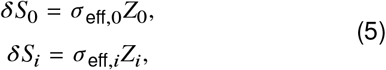

where *Z*_0_, *Z*_1_, …, *Z*_*K*_ are independent standard normal random variables. The same *Z*_0_ enters every shell estimate because all ADCs use the same averaged b = 0 reference, whereas *Z*_*i*_ is specific to shell *i*.

Linearising Equation 1 gives

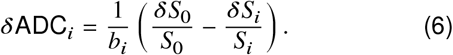

The marginal ADC uncertainty for shell *i* is

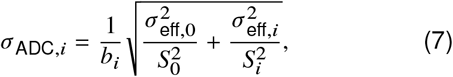

with *N*_0_ = 11 and *N*_*i*_ = 19–20 in our protocol. Using first-order error propagation (the delta method) (Oehlert, 1992), the temperature sensitivity and marginal uncertainty are

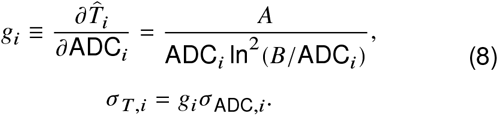

To account for the shared reference, we separate the temperature error into shared and shell-specific components with positive magnitudes

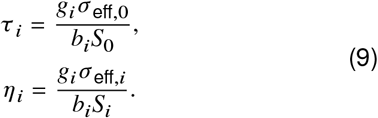

Let 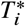 denote the temperature inferred from the noisefree b = 0 and shell-*i* mean signals. The first-order estimation error 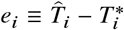 is

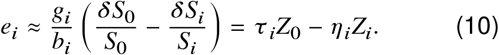

The shared reference induces positive cross-shell covariance. Changing the signs of the independent *Z*_*i*_ values leaves their joint distribution unchanged, so simulations may equivalently use *e*_*i*_ = *τ*_*i*_*Z*_0_ + *η*_*i*_*Z*_*i*_:

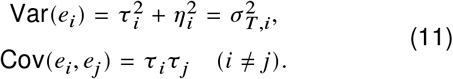

With ***e*** = (*e*_1_, …, *e*_*K*_)^T^ and ***τ*** = (*τ*_1_, …, *τ*_*K*_)^T^, the voxel-specific covariance matrix is

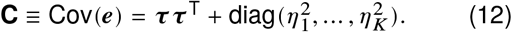

We define a per-voxel reference noise scale as the root-mean-square of the per-shell uncertainties:

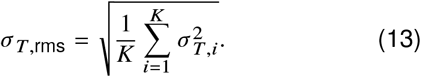

Applying the same positive normalisation to observed and simulated MAD leaves the classification unchanged.

### 2.3 Cross-shell consistency test

We assess cross-shell agreement by comparing the observed temperature spread with that expected from noise when the noise-free shell-wise temperatures 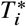 share a common value *T* ^*^. We quantify spread using the unscaled median absolute deviation (MAD), which limits the influence of one atypical shell:

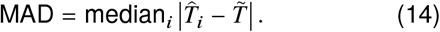

Observed and simulated MAD use the same unscaled definition. The null hypothesis is

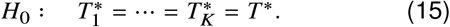

Under *H*_0_, 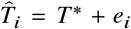, with errors given by Equation 10 and covariance by Equation 12. MAD is invariant to the common offset *T* ^*^, giving the noise-only value

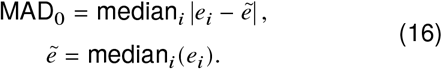

We define the dimensionless dispersion statistic

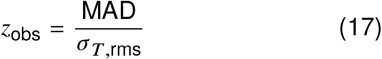

For each noise-only realisation *r* at voxel position **x**, we draw one shared *Z*_0_ and *K* shell-specific *Z*_*i*_ values using Equation 10 and calculate

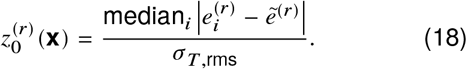

The median of this voxel-specific distribution, *z*_0.5_(**x**) = median_*r*_ 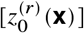, is the operating threshold. A voxel is classified as cross-shell consistent when

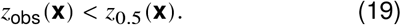

Thus, retained voxels have less dispersion than the median of their own noise-only distribution.

## 3 Methods

### 3.1 Participants and data acquisition

We analysed DWI data from 12 participants (seven female and five male; age range 58–79 years, mean 71.2 ± 5.9 years) drawn from a broader multimodal imaging study that included assessment of amyloid status. Six participants were classified as amyloid-positive and six as amyloid-negative using ^18^F-NAV4694 PET. Uptake was expressed on the Centiloid scale (Klunk et al., 2015), with positivity defined in this study as at least 20 Centiloids. The magnetic resonance imaging (MRI) session comprised a T1-weighted anatomical acquisition, task-based functional MRI, and then the multi-shell diffusion acquisition used here. All 12 participants had a complete diffusion acquisition and the accompanying opposite-phase-encoded reference images.

Ethical approval was obtained from the Human Research Ethics Committee of the University of Newcastle (H-2020-0439). All participants provided written informed consent.

Imaging was performed at the Hunter Medical Research Institute (HMRI) Imaging Centre on a 3 T Siemens MAGNETOM Prisma system (Siemens Healthineers, Erlangen, Germany; syngo MR E11) using a 64-channel head/neck coil. Multi-shell DWI used single-shot spin-echo echo-planar imaging with bipolar diffusion encoding. The acquisition comprised 11 b = 0 volumes, 20 diffusion directions at each of b = 200, 500, and 700 s/mm^2^, and 19 directions at each of b = 1100 and 1500 s/mm^2^. Sequence parameters were 2.0 mm isotropic native resolution, repetition time / echo time (TR / TE) = 2560 / 102 ms, flip angle = 90°, partial Fourier = 0.75, and multiband acceleration factor = 4. Images were acquired with a 108 × 108 matrix, 72 contiguous slices, anterior-to-posterior phase encoding, and a total readout time of 73.8 ms; the diffusion series lasted approximately 4 min 39 s. A pair of opposite-phase-encoded b = 0 spin-echo reference images was acquired for susceptibility-distortion correction. A T1-weighted magnetisation-prepared rapid gradientecho (MPRAGE) acquisition provided the anatomical reference at 0.9 mm isotropic resolution (TR / TE / inversion time = 1900 / 2.19 / 1000 ms; flip angle = 8°).

Infrared tympanic temperature was recorded to the nearest 0.1 °C in both ears using a Braun ThermoScan 7+ thermometer on entry to and exit from the MRI scanner room. Left and right measurements were averaged within each time point. The post-scanning mean was used as a descriptive external reference because it was temporally closer to the diffusion acquisition.

### 3.2 Preprocessing and candidate CSF mask

DWI data were preprocessed using MRtrix3 version 3.0.8 (Tournier et al., 2019). Magnitude-reconstructed DWI entered MP-PCA denoising (`dwidenoise`, Exp2 estimator, 5 × 5 × 5 patches), which provided the spatial input magnitude-noise map. Each denoised volume was corrected by numerical inversion of the Rician magnitude mean and estimated magnitude-noise standard deviation (Koay & Basser, 2006), followed by Gibbs-ringing correction using `mrdegibbs` (Kellner et al., 2016). Susceptibility distortion was estimated from the aligned opposite-phase-encoded reference images using FSL’s `topup` (Andersson et al., 2003). FSL’s `eddy` corrected the diffusion data for susceptibility distortion, eddy currents, and between-volume participant motion, with outlier-slice replacement (Andersson et al., 2016; Andersson & Sotiropoulos, 2016). Both tools (FSL 6.0.7.10) were run through `dwifslpreproc`. N4 bias-field correction (Tustison et al., 2010) used `dwibiascorrect ants` (ANTs 2.6.5). To reduce within-shell intensity drift, each volume, including b = 0 volumes, was rescaled to the median signal of its shell within the brain mask when the required adjustment was at least 0.5%. The resulting data were resampled from 2.0 to 1.5 mm isotropic resolution using cubic interpolation.

The MP-PCA noise map was divided by the saved N4 bias field before regridding, matching the spatial intensity scaling of DWI. The actual per-volume drift factors entered the mean-error coefficients through Equation 4. This transformed input-noise reference remains a first-order approximation: denoising, distortion correction, and interpolation can alter final residual variance and spatial dependence. Residual-bootstrap sensitivity analyses and a matched quadratic-correction control are reported in Supplementary Section S3.

The candidate CSF mask was derived from the base-line preprocessed data using MRtrix3’s multi-shell, multitissue constrained spherical deconvolution (`dwi2fod msmt_csd`) (Jeurissen et al., 2014). White matter, grey matter, and CSF response functions were estimated using `dwi2response dhollander`; the resulting tissue components were intensity-normalised using `mtnormalise`. The normalised CSF component was thresholded at 0.07. The binary mask was cleaned by one erosion–dilation cycle followed by one median-filtering pass. The same candidate mask and anatomical assignments were used for the primary analysis and matched quadratic-correction control. Temperatures, uncertainties, and consistency selection were calculated separately for each correction.

### 3.3 Analysis overview

Figure 1 summarises the analysis. Step 1, “Acquire multishell diffusion data”, supplies the b = 0 reference and four non-zero shells. In 2A, “Observed temperature dispersion”, each reference–shell pair yields a temperature estimate; the four estimates provide the voxel-wise median and cross-shell MAD. In parallel, 2B, “Expected dispersion from noise”, propagates the measured signal noise, including the shared-reference contribution, to a voxelspecific noise-only distribution. Step 3, “Compare each voxel with its own noise null”, retains voxels whose observed dispersion is below their own null median. Cluster-assisted anatomical assignment defines the lateral ventricular (LV) and subarachnoid (SA) compartments used for regional summaries.

**Figure 1.**
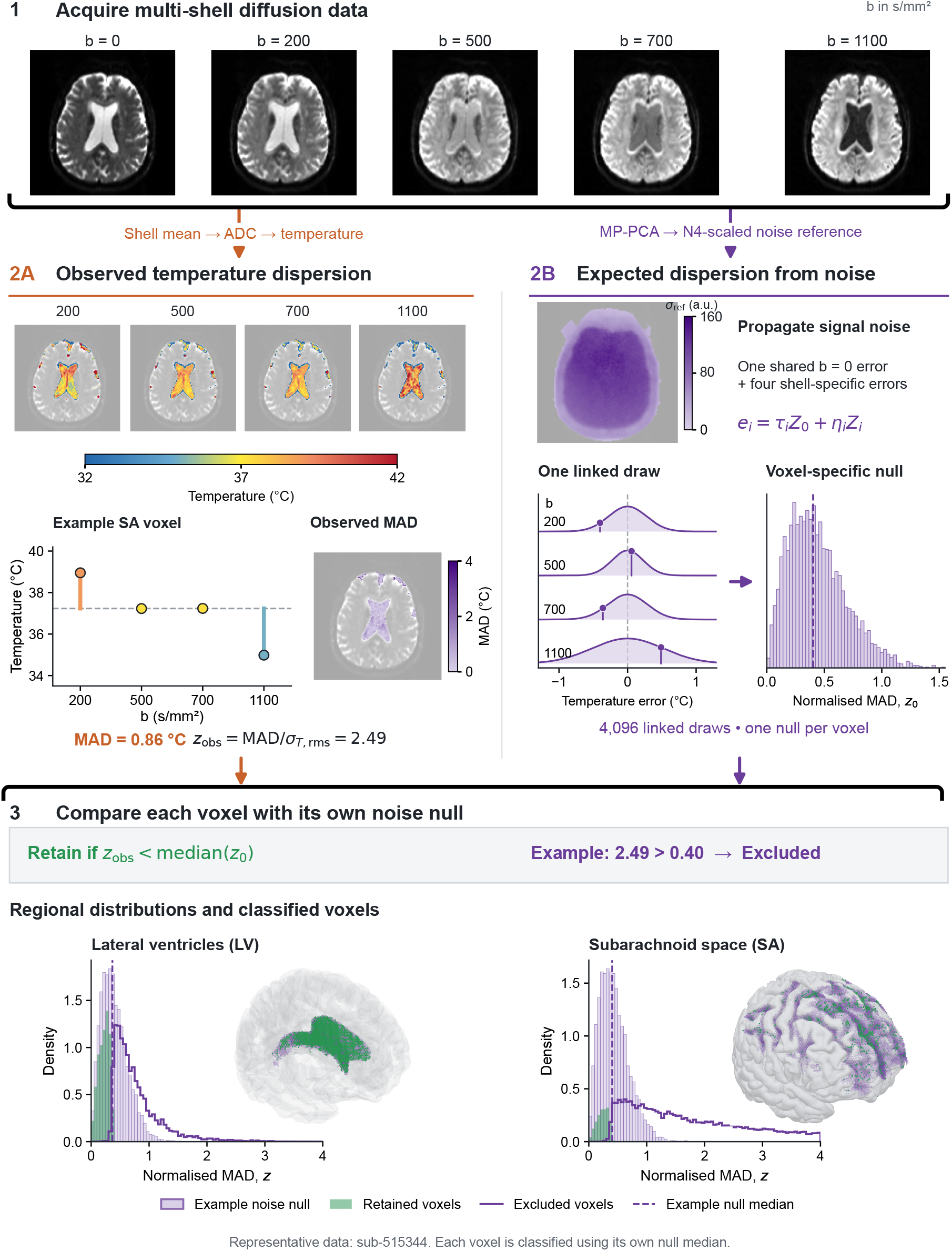
Voxel-wise multi-shell DWI thermometry and cross-shell consistency workflow. Representative data from sub-515344. (1) Multishell acquisition provides the b = 0 reference and four non-zero diffusion shells. (2A, Observed temperature dispersion) Each reference–shell pair is converted to an apparent diffusion coefficient (ADC) and then temperature using the Arrhenius relationship. The four temperatures yield the observed cross-shell median absolute deviation (MAD) and its normalised value, *z*_obs_. (2B, Expected dispersion from noise) Marchenko– Pastur principal component analysis (MP-PCA) provides the input-noise reference, scaled for the N4 bias field and actual volume drift when forming mean-error coefficients. Each linked draw combines one shared b = 0 error with four shell-specific errors, producing four correlated temperature errors and one noise-only *z*_0_ value. Repeating this 4,096 times produces the illustrated voxel-specific null distribution. (3, Compare each voxel with its own noise null) A voxel is retained when *z*_obs_ < median(*z*_0_). The example SA voxel is excluded because its observed temperature dispersion exceeds the median expected from noise (2.49 > 0.40). The lower panels show regional distributions and three-dimensional views of retained (green) and excluded (purple) voxels in the lateral ventricles (LV) and subarachnoid space (SA). The shaded noise distributions and dashed medians are illustrative examples; each voxel is classified using its own null median.

### 3.4 Shell-wise temperature estimation and multishell summary

For each candidate CSF voxel, the mean b = 0 signal and the mean signal within each non-zero shell were used to calculate ADC and temperature (Figure 1, 2A; Equations 1 and 2). We used the ACSF Arrhenius calibration (*A* = 2732 K, *B* = 19.649 mm^2^/s) (Shibukawa et al., 2021); sensitivity to the Mills water calibration is reported in Supplementary Section S1. The median across the four shells (b = 200, 500, 700, 1100 s/mm^2^), 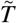, was the primary voxel-wise temperature estimate.

**Figure 2.**
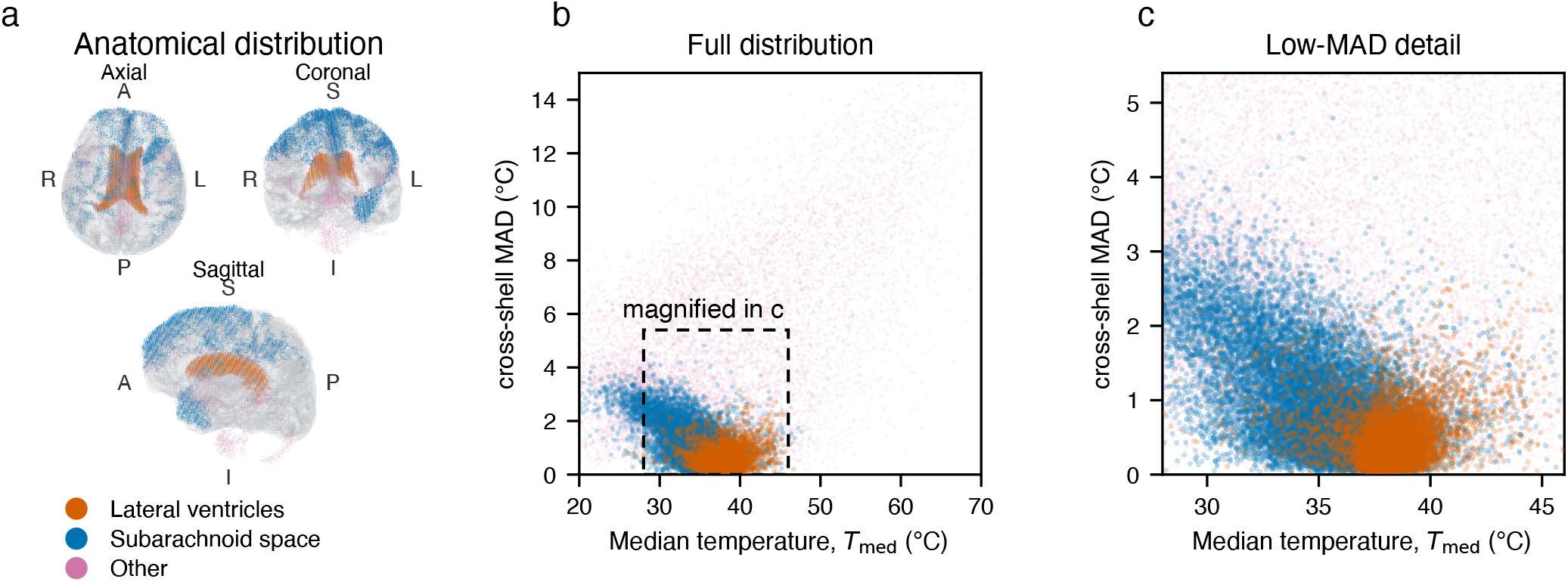
Spatial origin and cross-shell dispersion of candidate cerebrospinal fluid (CSF) voxels. Representative participant (sub-515344), using b = 200, 500, 700, and 1100 s/mm^2^, before consistency selection. (a) Axial, coronal, and sagittal orthographic whole-volume projections (up to 10,000 voxels per category) of lateral ventricles (LV; orange), subarachnoid space (SA; blue), and Other (pink), displayed with a transparent pial surface. Other denotes candidates not assigned to LV or SA. Orientation labels indicate anterior/posterior (A/P), right/left (R/L), and superior/inferior (S/I). (b) Per-voxel median temperature, 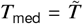, versus cross-shell median absolute deviation (MAD). (c) Enlargement of the square boxed region (temperature 28–46 °C; MAD 0–5.4 °C), with equal magnification of both axes. Scatter layering from front to back: LV, SA, Other.

Before assessing cross-shell consistency, we checked that each shell provided a usable temperature estimate. Reference and shell signals had to be finite and positive, ADC had to exceed 10^−6^ mm^2^/s, and temperature and its propagated uncertainty had to be finite. The uncertainty *σ*_*T,i*_ also had to be positive and below 20 °C. A voxel passed four-shell validity only if all four shells passed these checks.

The b = 1500 shell was excluded because of its greater residual sensitivity to the magnitude-noise floor (Supplementary Section S3). A matched sensitivity analysis recomputed temperature from the median of the b = 200, 500, and 700 estimates while retaining the voxel population defined by the four-shell analysis.

### 3.5 Noise propagation and consistency classification

In LV and SA, classification required four-shell validity and median temperature at least 25 °C. Per-shell uncertainties *σ*_*T,i*_ followed Equations 4–11. For each eligible voxel, scrambled Sobol quasi-Monte Carlo draws, generated using `scipy.stats.qmc.Sobol` (Joe & Kuo, 2008; Matoušek, 1998), sampled one shared b = 0 factor and four shell-specific factors (Figure 1, 2B). Each draw yielded a normalised noise-only MAD, *z*_0_ (Equation 18); its median across draws defined *z*_0.5_. All eligible voxels were initially evaluated with 2^12^ draws; those with |*z*_obs_ − *z*_0.5_| ≤ 0.015 were refined with 2^15^ draws. Additional independent scrambles and up to 2^21^ draws adjudicated boundary disagreements. Voxels were retained when their observed *z* was below their own null median (Figure 1, 3; Equation 19).

**Figure 3.**
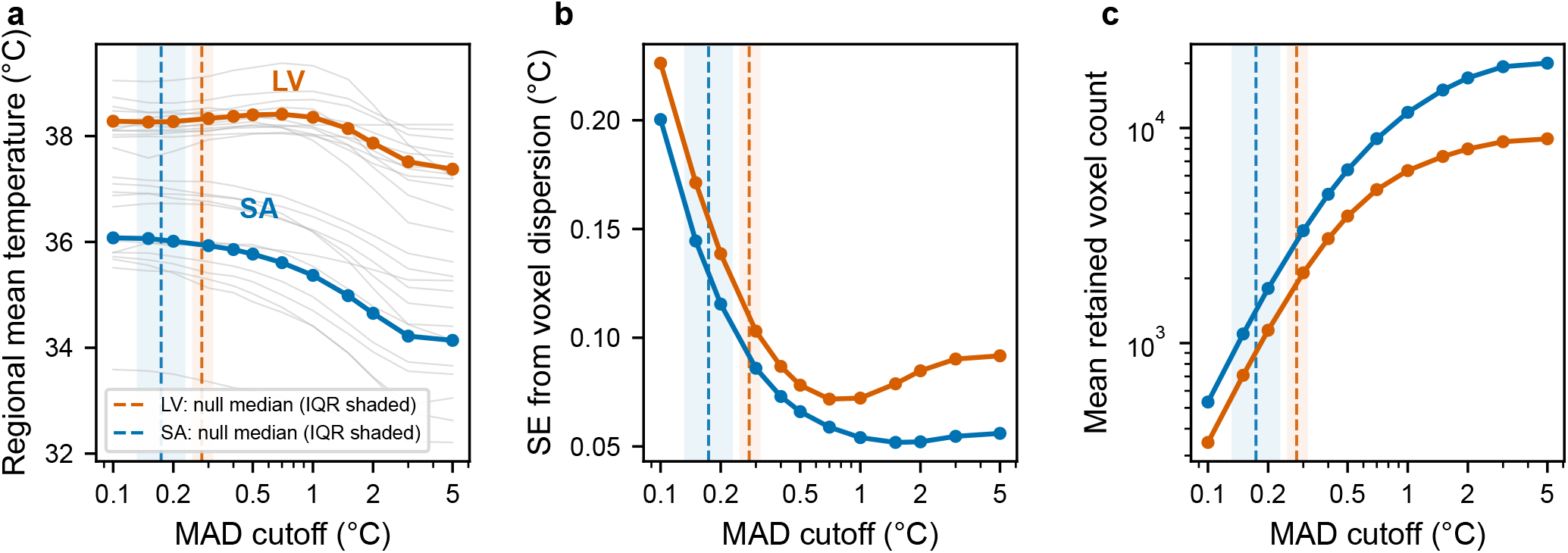
An exploratory absolute-MAD threshold sweep illustrates the trade-off between regional estimates, precision, and voxel retention. The cross-shell median absolute deviation (MAD) cutoff was swept from 0.1 to 5.0 °C in lateral ventricular (LV) and subarachnoid (SA) voxels across 12 participants. Dashed lines and shaded bands show pooled medians and interquartile ranges (IQRs) of the effective absolute thresholds associated with the voxel-specific noise nulls (medians 0.278 °C in LV and 0.174 °C in SA). (a) Regional mean temperature: grey lines show participants and coloured lines show group means. (b) Within-participant standard error of the regional mean, calculated from between-voxel temperature dispersion and corrected for spatial autocorrelation. (c) Mean retained voxel count.

Using the null median gives all voxels the same operating quantile while allowing its numerical value to depend on their measured noise. Under a correctly specified continuous null, approximately half of otherwise eligible noise-only voxels would pass this dispersion rule.

### 3.6 Cluster-assisted compartment definition and anatomical adjudication

Candidate CSF voxels satisfying the four-shell validity criteria in the baseline preprocessed data were partitioned using hierarchical density-based spatial clustering of applications with noise (HDBSCAN) (McInnes et al., 2017) on the feature vector [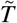, MAD, *x, y, z*]. Each feature was standardised to zero mean and unit variance within the participant. HDBSCAN used Euclidean distance, excess-of-mass cluster selection, `min_cluster_size = 500`, and `min_samples = 10`. Regional analyses applied each analysis-specific consistency selection within these fixed, anatomically adjudicated compartments.

All cluster assignments were manually adjudicated by anatomical location. Clusters corresponding to the lateral ventricles (LV) were labelled LV; clusters surrounding the pial surface were labelled as subarachnoid space (SA), with disconnected SA clusters pooled into one participant-level SA compartment; and remaining clusters were assigned to Other. Other also included inspected voxels carrying the HDBSCAN noise label (−1) and any remaining candidates not assigned to LV or SA. Regional temperature analyses used LV and SA voxels. A cluster fingerprint (centroid, spread, 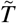, voxel count, and final anatomical assignment) was saved for reproducibility. FreeSurfer pial surfaces (Fischl, 2012) supported the anatomical identification of SA clusters and their display in Figure 2.

Temperature and MAD were clustering features, whereas accepted clusters were identified anatomically. The HDBSCAN noise label was recorded before consistency classification and described clustering membership, separately from cross-shell agreement.

### 3.7 Two-compartment tissue-admixture sensitivity analysis

To quantify how strongly conventional tissue admixture could bias the compartment estimates and their difference, we used a simplified two-compartment CSF–tissue signal model with canonical CSF and tissue diffusivities (Supplementary Section S2). For each participant and compartment, we calculated the b-dependent temperature contrast

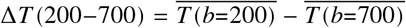

among voxels retained by the consistency criterion. The b = 200 and 700 shells were the lowest and highest of the three lower-b shells, maximising the predicted contrast from tissue admixture while avoiding the greater residual sensitivity of b = 1100 and 1500 to the magnitude-noise floor. The model was then used to determine how much tissue admixture would be required to reproduce the contrast (Supplementary Section S2). In that model, the CSF fraction refers to its contribution to the b = 0 signal, rather than an anatomical volume fraction. By attributing the entire positive contrast to tissue, we calculated an implied signal fraction and the corresponding downward bias of the four-shell median. The resulting values were sensitivity bounds, quantifying the effect of b-dependent tissue admixture under the stated model.

### 3.8 Single-shell and multi-shell agreement

We compared the primary multi-shell analysis with conventional single-shell DWI thermometry using the derivative-bounded histogram method separately at b = 200, 500, 700, and 1100 s/mm^2^ within each anatomically assigned compartment (Sakai et al., 2011; Sakai et al., 2012). Single-shell histograms used anatomically assigned candidates with finite temperatures between 25 and 45 °C. Each histogram had 256 bins, and an eighth-order polynomial was fitted over the full range. The dominant fitted maximum was identified, and the two flanking inflection points defined *T*_low_ and *T*_high_, corresponding to the positive and negative extrema of the first derivative. Following the method of Sakai et al., the single-shell temperature was the mean of the voxel temperatures within these bounds. The multi-shell estimate was the regional mean of the within-voxel four-shell medians after consistency selection. For each participant, compartment, and shell, the difference was *T*_single,Sakai_(*b*) − *T*_multi,mean_.

Additional diagnostics were calculated from the same single-shell temperatures within the 25–45 °C range: the mean and median before applying the Sakai bounds, the fraction within those bounds, and the fraction of those voxels also retained by the multi-shell criterion. These summaries characterised the effect of voxel selection.

### 3.9 Statistical analysis

For each participant and compartment, the regional temperature was the mean of 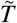 across voxels passing the consistency criterion (Figure 1, 3). To estimate its conditional input-reference noise uncertainty, we perturbed the observed shell temperatures using the linked error model (Equation 10) and recalculated the within-voxel median for 16,384 scrambled Sobol draws. The standard deviation of these simulated medians supplied 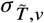 for each retained voxel *v*, separately from the RMS shell scale used for classification. Voxel selection was held fixed. These uncertainties were combined as their regional rootmean-square. To account approximately for spatial dependence, we used the nearest-neighbour model with *ρ*_nn_ ≈ 0.64 and approximately three undirected pairs per voxel, giving a variance-inflation factor of 4.84 and *N*_eff_ = *N*/4.84. The regional noise standard error was the regional root-mean-square median uncertainty divided by 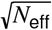 (Vallejos & Osorio, 2014). These estimates quantify random measurement-noise uncertainty conditional on the retained voxel population and the stated spatialcorrelation model (Supplementary Section S2). Group regional values are reported as the mean ± sample standard deviation (SD) of the 12 participant-level estimates.

The LV–SA temperature difference was computed for each participant and tested against zero using a twosided paired t-test, with a 95% confidence interval (CI) for the mean paired difference. The consistency of its direction across participants was additionally assessed with an exact two-sided sign test. As an exploratory internal comparator, absolute cross-shell MAD cutoffs from 0.1 to 5.0 °C were applied to four-shell-valid LV and SA candidates with median temperatures of 25–80 °C. At each cut-off, we recorded the regional mean, retained voxel count, and standard error based on the observed between-voxel temperature dispersion with the same spatial-correlation correction. These empirical standard errors differ from the propagation-based measurement-noise scales defined above. All remaining association analyses were exploratory. The association between retained SA voxel count and SA temperature was assessed using Spearman correlation with a two-sided p-value from a fixed-seed Monte Carlo permutation test with 999,999 permutations. The largest half of the cohort by retained SA compartment size (*n* = 6) was examined in a post hoc descriptive sensitivity analysis. Post-scanning tympanic temperature was used as a descriptive external reference measurement. The association between retained SA voxel count and the absolute SA–ear difference was assessed using Spearman correlation with the same permutation procedure; agreement in these six participants was summarised by the paired SA–ear difference and its 95% confidence interval. Exploratory p-values were reported unadjusted.

## 4 Results

### 4.1 Candidate CSF compartments and cross-shell dispersion

The fixed anatomical assignments identified LV and at least one SA cluster in every participant. Four-shell validity checks (Section 3.4) were satisfied by 109,302 LV and 248,396 SA candidates across the cohort. A further 234,616 valid candidates not assigned to either compartment were categorised as Other, including remaining clusters and unassigned voxels. Only LV and SA voxels entered the regional analyses. The consistency criterion retained 22,476 LV and 18,367 SA voxels. Participant-level retained counts ranged from 598 to 7,171 in LV (median 1,249.5) and from 295 to 3,852 in SA (median 1,076).

Figure 2 shows the signal-valid candidate voxels in a representative participant before consistency selection: LV in orange, SA in blue, and Other in pink.

In this participant, LV and SA voxels predominantly occupied the low-MAD range, with median cross-shell MADs of 0.37 °C and 0.76 °C, respectively. Other voxels had a broader distribution and a median MAD of 3.31 °C, dominating the high-dispersion tail (Figure 2b). All three categories nevertheless overlapped in the low-MAD range (Figure 2c). Consistency selection compared each voxel’s dispersion with that expected from its propagated measurement noise.

### 4.2 Noise propagation provides a voxel-specific consistency criterion

The primary consistency criterion used each voxel’s propagated noise distribution and was defined independently of the observed temperature distribution. A voxel was retained when its normalised cross-shell MAD was below the median of its own noise-only distribution. The pooled median noise-null threshold was *z* = 0.373 in LV and *z* = 0.409 in SA; the pooled median absolute MAD thresholds were 0.278 °C and 0.174 °C, respectively. These values summarise voxel-specific thresholds; selection used each voxel’s own threshold. After spatial-correlation correction, the conditional regional input-reference noise standard error was below 0.05 °C in LV and 0.04 °C in SA.

As an exploratory comparator, we swept absolute MAD cutoffs from 0.1 to 5.0 °C in both compartments across all 12 participants, retaining voxels with MAD below each cutoff. We tracked regional mean temperature, the spatial-correlation-corrected standard error calculated from between-voxel temperature dispersion, and retained voxel count (Figure 3). These empirical standard errors are distinct from the propagation-based measurement-noise values above.

Across absolute MAD cutoffs from 0.1 to 1.0 °C, the LV group mean ranged from 38.27 to 38.41 °C, before decreasing to 37.37 °C at a cutoff of 5.0 °C. The SA mean decreased from 36.08 °C at 0.1 °C to 35.37 °C at 1.0 °C and 34.14 °C at 5.0 °C as colder, high-MAD voxels were included (Figure 3a). At the strictest cut-off, the SA mean was 0.03 °C above the primary noise-referenced estimate. The empirical standard-error minima occurred near cutoffs of 0.7 °C in LV and 1.5 °C in SA (Figure 3b). At the SA minimum, mean temperature was 34.98 °C, 1.07 °C below the primary estimate. Retained voxel count increased monotonically (Figure 3c). The SA standard-error minimum thus occurred at a colder regional estimate than the primary criterion.

### 4.3 LV temperature estimates were higher than SA estimates

Among voxels passing the consistency criterion, mean LV temperature was 38.41 ± 0.37 °C and mean SA temperature was 36.05 ± 1.10 °C using the ACSF calibration. The paired within-participant LV–SA difference was 2.36 ± 1.17 °C (95% CI 1.61–3.10 °C; paired *t*(11) = 6.98, *p* < 0.001) and was positive in all 12 participants (exact two-sided sign test, *p* = 0.000488; Figure 4).

**Figure 4.**
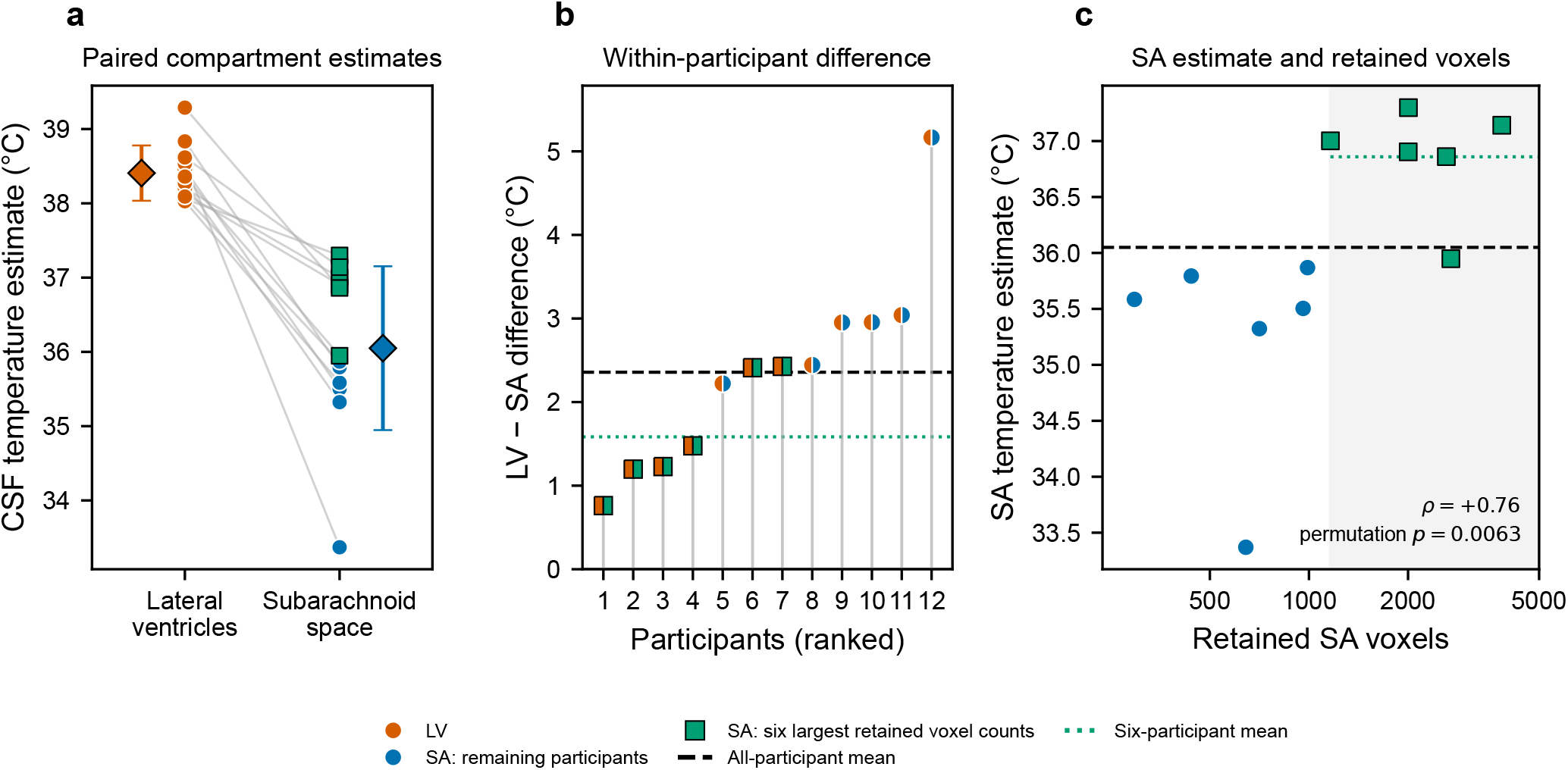
Lateral ventricular (LV) and subarachnoid (SA) temperature estimates and association with retained SA voxel count. Orange denotes LV and blue denotes SA; green, black-outlined squares identify SA measurements from the six participants with the largest retained SA voxel counts. Black dashed lines show all-participant means and green dotted lines show means for this subset. (a) Paired LV and SA estimates; diamonds and error bars show group means ± SD. (b) Within-participant LV–SA differences ranked by magnitude. Each split marker represents one participant: the orange left half denotes LV and the blue or green right half denotes SA. Outlined squares identify the same six-participant subset. (c) SA temperature versus retained SA voxel count. Shading identifies the post hoc subset of the six participants with at least 1,162 retained voxels.

Across all 12 participants, the SA estimate increased with retained SA voxel count (Spearman *ρ* = +0.76, permutation *p* = 0.0063; Figure 4c). In a post hoc subset of the six participants with the largest retained SA voxel counts, each containing at least 1,162 voxels, mean SA temperature was 36.86 ± 0.47 °C and the mean LV–SA difference was 1.58 ± 0.69 °C. Within this subset, the correlation between retained count and SA temperature was *ρ* = −0.14.

Compared with post-scanning tympanic temperature, the mean SA–ear temperature difference was −0.7 ± 1.2 °C across all 12 participants. The absolute discrepancy decreased as retained SA voxel count increased (*ρ* = −0.66, permutation *p* = 0.024). In the six participants with the largest retained SA voxel counts, mean postscanning tympanic temperature was 36.8 ± 0.1 °C and the paired SA–ear difference was +0.1 ± 0.5 °C (95% CI −0.5 to +0.6 °C; paired t-test *p* = 0.778; Supplementary Figure S1).

### 4.4 Sensitivity to tissue admixture

The endpoint temperature contrast Δ*T*(200−700) among retained voxels was 0.371 ± 0.224 °C in LV and 0.245 ± 0.347 °C in SA. The paired LV-minus-SA endpoint-contrast difference was 0.126 ± 0.214 °C (95% CI −0.011 to 0.262 °C). Attributing each compartment’s contrast entirely to tissue admixture under the model in Section 3.7 implied CSF b = 0 signal fractions of 0.988 in LV and 0.992 in SA, with predicted downward biases of 0.68 and 0.45 °C in the four-shell median. Conversely, a modelled CSF signal fraction of 0.95 produced a 2.84 °C downward bias, comparable to the observed 2.36 °C regional difference, but predicted Δ*T*(200−700) = 1.45 °C—5.9-fold larger than the observed mean SA contrast. Under the assumed canonical CSF–tissue diffusivities, tissue admixture sufficient to reproduce the regional level difference was incompatible with the weak observed b-dependence.

Adding the predicted biases to both compartment estimates changed the mean paired difference from 2.36 to 2.59 °C. Adding the predicted bias only to SA left a difference of 1.91 °C. The modelled biases therefore partly cancelled in the paired difference, while the LV–SA ordering persisted in both sensitivity scenarios.

### 4.5 Sensitivity of the temperature summary to b = 1100

Omitting b = 1100 from the temperature summary had little effect on regional estimates within the population selected by the four-shell criterion. Recalculating the within-voxel median using b = 200, 500, and 700 yielded 38.37 ± 0.38 °C in LV, 36.07 ± 1.09 °C in SA, and a paired difference of 2.31 ± 1.16 °C. Relative to the primary four-shell summary, the cohort means changed by −0.03 °C in LV, +0.02 °C in SA, and −0.05 °C for the paired difference. This analysis held consistency classification and compartment assignment fixed. Sensitivity to the common b = 0 reference is reported in Supplementary Section S3.

### 4.6 Comparison with conventional single-shell DWI thermometry

As an implementation check, the single-shell LV temperature estimated from the b = 0 and b = 700 s/mm^2^ images using the method of Sakai et al. was 37.5 ± 0.4 °C with the Mills calibration. Detailed calibration and estimator checks are reported in Supplementary Section S1.

We then applied the conventional estimator independently at b = 200, 500, 700, and 1100 s/mm^2^ using the same ACSF calibration as the primary analysis. The cohort mean ± SD conventional temperatures were, respectively, 38.71 ± 0.51, 38.35 ± 0.26, 38.52 ± 0.31, and 39.86 ± 0.65 °C in LV, and 36.10 ± 1.24, 35.22 ± 1.27, 34.98 ± 1.34, and 33.97 ± 1.56 °C in SA. We compared these conventional single-shell estimates with the primary multi-shell estimates (Figure 5). The comparison reflects differences in both voxel selection and regional summarisation. In LV, the median single-shell minus multi-shell differences were +0.32, −0.01, +0.10, and +1.43 °C at b = 200, 500, 700, and 1100 s/mm^2^, respectively. Differences were smaller at b = 200–700 than at b = 1100, which produced a higher estimate in every participant.

**Figure 5.**
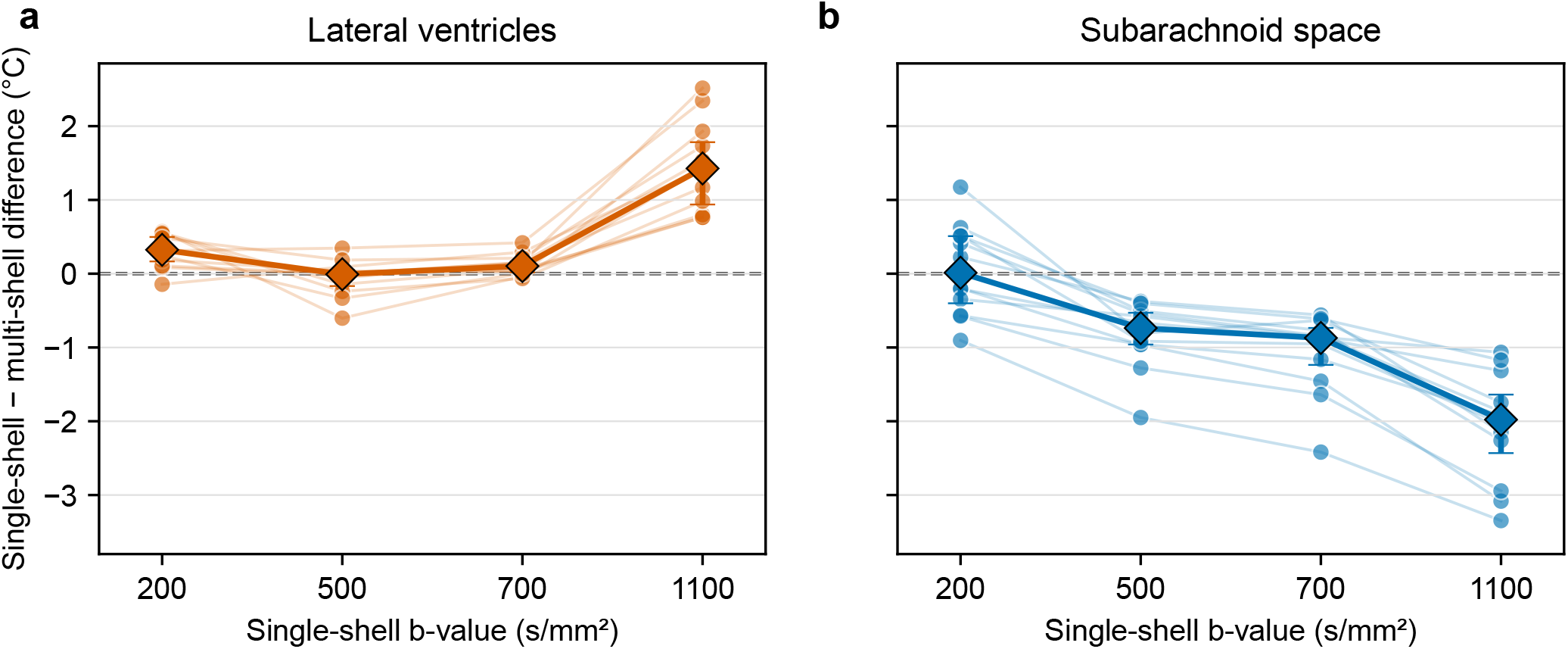
Conventional single-shell estimates differ from the primary multi-shell estimate by compartment and b-value. The plotted difference is the Sakai derivative-bounded mean minus the primary regional multi-shell estimate; both use the ACSF calibration. The con-ventional estimate used anatomically assigned candidates with finite single-shell temperatures in the 25–45 °C histogram range, whereas the primary estimate used the four-shell within-voxel median among voxels passing the consistency criterion. The comparison therefore includes differences in both voxel selection and regional summarisation. (a) Lateral ventricles (LV; orange). (b) Subarachnoid space (SA; blue). Faint lines and circles show the 12 individual participants; diamonds and error bars show the cohort median and interquartile range. The dashed line denotes agreement. In SA, single-shell estimates were lower in every participant at b = 500–1100 s/mm^2^.

In SA, the corresponding median differences were +0.01, −0.74, −0.87, and −1.98 °C. Although b = 200 gave a small median temperature difference, a median of only 9.5% of voxels within the Sakai averaging window passed the consistency criterion. At b = 500, 700, and 1100, the single-shell estimate was lower than the primary multishell estimate in every participant. No single b-value therefore showed the same agreement across both compartments.

## 5 Discussion

### 5.1 Principal findings

Multi-shell DWI provided a noise-referenced internal consistency check for regional CSF thermometry. Temperature estimates were higher in LV than in SA in all 12 participants, with a mean paired difference of 2.36 ± 1.17 °C. This ordering persisted across the reported sensitivity analyses. SA temperature was associated with retained SA voxel count, and the magnitude of the LV–SA difference was smaller in the six participants with the largest counts. Regional sampling is therefore important to the interpretation of these estimates.

Multi-shell analysis contributed differently in the two compartments. Conventional single-shell LV estimates at b = 200–700 s/mm^2^ closely agreed with the primary estimate, while SA estimates varied more with b-value and voxel selection. No single b-value showed the same agreement in both compartments. Multi-shell measurements provided a consistency assessment beyond the ventricles and supported separate regional estimates.

### 5.2 What multi-shell analysis adds

The consistency criterion evaluates each voxel’s temperature disagreement against its own noise-only distribution, including covariance from the shared b = 0 reference. Absolute MAD cutoffs depend on signal level, image noise, b-values, and calibration. The exploratory sweep showed how changing the cutoff affected regional temperature, retained count, and empirical standard error (Figure 3).

Relaxing the absolute cutoff increased retained SA count and made the regional mean colder, even as the empirical standard error decreased over part of the sweep. Across participants, however, larger SA populations retained by the noise-referenced criterion were associated with warmer estimates. Retained count therefore needs to be interpreted together with the selection criterion.

Classification retains measurements with dispersion below their own noise-null median. Regional precision depends on averaging and spatial correlation. Cross-shell agreement provides an internal consistency check; shell-independent biases require additional assessment.

Multi-shell data also provided the endpoint temperature contrast Δ*T*(200−700) for the supporting tissueadmixture sensitivity analysis. The simplified model tests one plausible alternative explanation for the compartment difference and shows whether its predicted bias is shared by the two compartments.

### 5.3 Subarachnoid sampling and regional measurement quality

Larger retained SA populations were associated with warmer SA estimates across the 12 participants. The six participants with the largest counts also had lower between-participant variability and a smaller, still positive LV–SA difference. Mild residual tissue admixture is one plausible contributor: it can lower apparent diffusivity while preserving cross-shell consistency. Retained count also depends on anatomy, noise, clustering, and the consistency criterion, so the association may reflect several contributing factors. The restricted count range and small subset limit conclusions about a sampling plateau.

Across participants, the absolute difference between SA and post-scanning tympanic temperature decreased as retained SA voxel count increased. In the six participants with the largest counts, the paired SA–ear difference was +0.1 ± 0.5 °C (Supplementary Figure S1 and Supplementary Table S9), providing an external plausibility check for the SA estimates.

The minimum retained count in this post hoc subset was 1,162 voxels. Raw counts depend on acquisition and reconstruction resolution, smoothing, and spatial correlation. Regional estimates should therefore be accompanied by retained voxel count, effective sample size, and uncertainty from propagated measurement noise.

### 5.4 Choice of diffusion weightings

The included shells provide complementary information. At b = 200, relatively high CSF signal supports temperature estimation; b = 500 and 700 add diffusion weightings for assessing agreement and the endpoint contrast used in the tissue-admixture model. The b = 1100 shell extends the consistency assessment at the lowest SNR of the included shells. We retained it in the primary analysis and tested its influence on the temperature summary separately, holding the selected population fixed. The b = 1500 shell was excluded because it was more sensitive to the magnitude-noise floor. These analyses used an existing acquisition. Prospective thermometry protocols should evaluate the balance between diffusion sensitivity, motion compensation, SNR, and scan time, including the number of shells needed for consistency assessment.

### 5.5 Relationship to previous DWI thermometry

In the present LV analysis, single-shell estimates at b = 200–700 s/mm^2^ were closer to the primary multi-shell estimate than the b = 1100 estimates (Figure 5). With the Mills calibration, the b = 700 implementation estimate of 37.5 ± 0.4 °C was consistent with previous singleshell DWI studies, which have generally reported healthy-cohort LV temperature estimates around 37–39 °C (Asfuroğlu et al., 2022; Horiuchi et al., 2023; Kozak et al., 2010; Sakai et al., 2012; Sakai et al., 2022; Sumida et al., 2016). The conventional histogram estimator and primary regional mean gave broadly similar LV estimates despite their different summaries and voxel selections.

The SA comparison was different. At b = 200 s/mm^2^, the conventional estimate showed a small median difference from the primary estimate, although a median of only 9.5% of voxels within its averaging window passed the consistency criterion. At b = 500–1100, the conventional estimate was lower in every participant. Shibukawa et al. (2021) used a b = 400 s/mm^2^ protocol with second-order motion-compensated gradients and fractional anisotropy (FA) < 0.1 filtering. FA filtering can reject directional abnormalities while retaining approximately isotropic tissue admixture. Such mixtures can produce b-dependent apparent diffusivity, giving cross-shell consistency sensitivity to contamination that a directional filter may miss.

Motion-compensated acquisition and subsequent cross-shell consistency assessment address different aspects of motion sensitivity. Second-order motion compensation reduces sensitivity to specified motion components during acquisition (Shibukawa et al., 2021; Stoeck et al., 2016). Cross-shell assessment evaluates remaining disagreement after acquisition. Here, elevated apparent temperatures along ventricular outflow pathways and around the brainstem coincided with high MAD and were excluded from regional estimates, consistent with b-dependent motion effects. Combining the two approaches could increase usable measurements in flow-prone regions while assessing residual cross-shell disagreement; this potential gain requires direct evaluation. The post-processing comparison, filter overlap, and acquisition details are reported in Supplementary Section S5 and Supplementary Figure S2.

Using cardiac-timed acquisitions and a thresholded ventricular estimator in 104 healthy volunteers, Sakai et al. (2022) found no significant effect of the tested CSF flow conditions on the DWI–MRS temperature difference, or association of this difference with aqueduct flow speed.

### 5.6 Physiological interpretation of the LV–SA gradient

The measured endpoint contrasts constrained the tissueadmixture explanation within the specified CSF–tissue model. Attributing the LV and SA contrasts entirely to tissue predicted downward biases of 0.68 and 0.45 °C, respectively. These biases partly cancelled in the paired difference, changing it from 2.36 to 2.59 °C; an SA-only scenario left 1.91 °C. Tissue admixture sufficient to explain the full regional difference predicted much stronger b-dependence than observed under the assumed diffusivities.

The higher ventricular than subarachnoid estimate is compatible with spatial variation in intracranial temperature. Magnetic resonance spectroscopy found a mean maximal spatial temperature range of 2.4 °C in brain tissue, with the highest values in the thalamus (Rzechorzek et al., 2022). These findings provide physiological context for the regional CSF estimates. Separate regional CSF estimates could help investigate heat exchange among CSF, brain tissue, blood, and the environment.

### 5.7 Limitations

Four considerations affect absolute temperature accuracy, interpretation of the regional difference, and generalisability.

1. **Residual flow**. CSF motion can increase ADC and apparent temperature. A flow-related ADC offset shared across shells would remain undetected by the consistency criterion and could differ between LV and SA. Residual flow therefore remains the main uncertainty in interpreting the full LV–SA difference as a physiological temperature difference. Comparisons with motion-compensated acquisition would help assess its contribution.
2. **Absolute scale and signal model**. Shibukawa et al. (2021) applied the ACSF relation as a secondary correction to human estimates obtained with the Mills water relation. Both calibrations preserved the LV– SA ordering in our data while changing absolute values and the difference magnitude (Supplementary Section S1). Protocol-matched phantom measurements are needed to quantify acquisition and calibration bias, while interpretation of the in vivo difference requires assessment of residual motion. Mild bdependence and magnitude-noise sensitivity indicate that the mono-exponential model is approximate, particularly at higher diffusion weightings (Supplementary Section S3). With four shells, the voxel-level criterion also has limited resolution for distinguishing patterns of disagreement. CSF composition is a further potential source of calibration uncertainty. Previous work found no significant association between amyloid PET uptake and DWI-derived ventricular temperature in either Alzheimer’s disease or control groups (Matsushita et al., 2024). A composition-related change in diffusivity that preserves mono-exponential attenuation would not be detected by cross-shell consistency; its effect on the LV–SA difference would depend on whether the bias differed between compartments.
3. **Subarachnoid sampling and cohort scope**. Age-related changes in extraventricular CSF volume may affect regional sampling and limit generalisation from this older cohort (Matsumae et al., 1996). Larger co-horts and repeat acquisitions are needed to quantify between-participant variation and test–retest repeatability. The six-participant subset and tympanic comparisons were post hoc; prospective sampling criteria require evaluation in an independent cohort.
4. **Semi-automated compartment definition**. Temperature and MAD were HDBSCAN features, so they could affect cluster membership and assignment to the noise label. The CSF mask was defined from diffusion tissue components before clustering, and accepted clusters were identified as LV or SA anatomically. Saved cluster fingerprints make these assignments auditable; comparison with spatial-only or independently segmented assignments would provide an additional validation.

### 5.8 Outlook

Separate ventricular and subarachnoid temperature estimates could be used to test whether central and superficial CSF respond differently to physiological changes or disease. The framework supports reporting regional temperatures together with retained voxel counts, effective sample sizes, and uncertainty from measurement noise. Prospective studies with motion-compensated diffusion encoding, protocol-matched phantoms, and repeat acquisitions are needed to assess accuracy, repeatability, and participant-level sampling criteria before diagnostic or monitoring applications can be evaluated.

## 6 Conclusion

We developed a multi-shell approach to regional CSF thermometry that uses measured image noise to assess agreement among shell-wise temperature estimates. In 12 participants, regional analysis identified higher diffusion-derived temperature in LV than in SA in every participant, with a mean paired difference of 2.36 °C. This ordering persisted across the reported sensitivity analyses, while the absolute SA estimate was associated with retained voxel count. Independent validation is needed to establish absolute accuracy and distinguish physiological temperature differences from residual motion effects. The framework provides an internal consistency check for separate ventricular and subarachnoid CSF thermometry and a basis for evaluating future acquisitions.

## 7 Data and Code Availability

A versioned reproducibility package containing the multi-shell temperature estimator, voxel-specific shared-reference consistency criterion, analysis settings, and a fully synthetic BIDS-compatible acquisition with reference outputs is publicly available in release v1.1.1 (Supplementary Section S6 and Figure S3). The source code is available in the public repository. The synthetic data contain no participant-derived images, metadata, anatomical templates, or fitted participant parameters. The package provides the numerical core and synthetic reference outputs for the primary calculation.

Participant MRI and participant-level derived datasets are not publicly available. Researchers may contact the corresponding author to request access. Requests will be assessed against participant consent and applicable ethics and institutional governance requirements; any sharing would be subject to the necessary approvals and data-sharing arrangements.

## 8 Author Contributions

Christian Behler: Conceptualization; Data curation; Formal analysis; Funding acquisition; Investigation; Methodology; Software (subsequent pipeline development); Validation; Visualization; Writing – original draft; Writing – review and editing. Michael Breakspear: Funding acquisition; Project administration; Resources; Supervision; Writing – review and editing. Saadallah Ramadan: Project administration; Resources; Supervision.

Thuy Thanh Dao: Data curation; Software (initial pipeline); Writing – review and editing. Thomas G. Close: Software. Sjoerd B. Vos: Methodology; Writing – review and editing, including scientific advice informing the Supplementary Material. Renate Thienel: Investigation; Project administration; Writing – review and editing.

Joseph Giorgio: Funding acquisition; Investigation; Project administration; Writing – review and editing. Anna Behler: Conceptualization; Data curation; Funding acquisition; Investigation; Methodology; Project administration; Resources; Software; Writing – review and editing. Michael Fay: Project administration; Resources; Supervision.

## 9 Funding

Data acquisition for this study was funded by the Hunter Medical Research Institute Brain Neuromodulation Research Program.

This work was supported by the Mark Hughes Foundation Centre for Brain Cancer Research through the Garry Willgoose PhD Scholarship, and by Tour de Cure Australia through a PhD Support Scholarship, both awarded to Christian Behler.

Michael Breakspear was supported by the National Health and Medical Research Council (APP2008612 and APP1152623).

## 10 Declaration of Competing Interests

No competing interests have been reported to the corresponding author.

## 11 Use of Generative Artificial Intelligence

Generative artificial intelligence tools (Claude, Anthropic; Codex, OpenAI; and Gemini, Google) were used during manuscript preparation to assist with language editing, critical review, and code development. The authors retain responsibility for the analyses, numerical outputs, interpretations, and submitted text.

## 12 Acknowledgements

We acknowledge the Traditional Custodians of the lands on which this research was conducted and developed, including the Awabakal and Worimi peoples, the Turrbal and Jagera peoples, the Gadigal people of the Eora Nation and the Whadjuk Noongar people. We recognize that sovereignty over these lands was never ceded and pay our respects to Elders past and present.

We also acknowledge the facilities and scientific and technical assistance of the National Imaging Facility (NIF), a National Collaborative Research Infrastructure Strategy (NCRIS) capability, at the Hunter Medical Re-search Institute Imaging Centre, University of Newcastle, and the Western Australia NIF node at the University of Western Australia.

## Notes

https://github.com/CB9683/multi-shell-dwi-csf-thermometry/releases/tag/v1.1.1

### Competing Interest Statement

The authors have declared no competing interest.

