## Supplement for "Noise-referenced multi-shell diffusion-weighted imaging for regional cerebrospinal fluid thermometry"

[†]Joint senior authors

#### S1. Calibration and conventional-estimator validation

Absolute temperature was calculated from the apparent diffusion coefficient (ADC) as

$$T = \frac{A}{\ln(B/\mathrm{ADC})} - 273.15.$$

Published CSF DWI thermometry has generally traced the ADC–temperature conversion to the pure-water diffusion measurements reported by Mills (1973). This relation was used in the foundational ventricular DWI thermometry studies (Kozak et al., 2010; Sakai et al., 2011; Sakai et al., 2012; Sumida et al., 2016; Tofts et al., 2008). Shibukawa et al. (2021) likewise generated the primary human temperature maps with the Mills relation. Their phantom experiment then showed that ACSF produced lower Mills-calibrated temperatures than water, and they derived an ACSF-specific relation as a secondary correction; in their preferred acquisition and filter combination, this correction changed the reported mean from 35.62 °C to 37.05 °C.

We therefore used the published ACSF relation of Shibukawa et al. (2021) for the primary analysis, with $A = 2732$ K and $B = 19.649$ mm$^2$/s, and treated the conventional Mills relation as a sensitivity analysis. The Mills calibration uses $A = 2256.74$ K and $B = 4.39221$ mm$^2$/s. We converted each retained voxel from its ACSF-calibrated temperature back to ADC and then applied the Mills constants. The calibration changed the absolute temperatures and the numerical difference, while preserving the ordering between the lateral ventricles (LV) and subarachnoid space (SA; Table S1).

Table S1: Calibration sensitivity of regional temperature and the paired difference. Values are group mean ± SD across 12 participants.

| Quantity | ACSF calibration (°C) | Mills calibration (°C) |
|---|---|---|
| Lateral ventricles | 38.41 ± 0.37 | 37.24 ± 0.45 |
| Subarachnoid space | 36.05 ± 1.10 | 34.42 ± 1.32 |
| Paired LV–SA difference | 2.36 ± 1.17 | 2.83 ± 1.40 |

As a conventional-estimator check, the derivative-bounded histogram estimate of Sakai et al. at b = 700 s/mm$^2$ under the Mills calibration was 37.47 ± 0.42 °C in LV (Sakai et al., 2011; Sakai et al., 2012), compared with 37.24 ± 0.45 °C for the Mills-calibrated multi-shell estimate. The derivative-bounded estimate was consistent with previous single-shell DWI reports (Kozak et al., 2010; Sakai et al., 2022; Sumida et al., 2016). Main Figure 5 uses the same derivative-bounded histogram method, with the ACSF calibration applied to both the single-shell and multi-shell estimates.

### S2. Partial-volume and spatial-correlation sensitivity

The two-compartment sensitivity analysis asked whether conventional tissue admixture, represented using canonical CSF and tissue diffusivities, could account for the observed LV–SA temperature difference. Its fraction parameter describes the contribution of cerebrospinal fluid (CSF) to the b = 0 signal. The model signal was

$$S(b; f) = S(0)\left[f \exp(-bD_{\mathrm{CSF}}) + (1-f)\exp(-bD_{\mathrm{tissue}})\right],$$

where $f$ is the fraction of the b = 0 signal attributed to CSF. It is equivalent to an anatomical volume fraction only under the additional assumption that CSF and tissue have equal b = 0 signal per unit volume. We used $D_{\mathrm{CSF}} = 3.0 \times 10^{-3}$ mm$^2$/s and $D_{\mathrm{tissue}} = 0.7 \times 10^{-3}$ mm$^2$/s (Holz et al., 2000; Pierpaoli et al., 1996; Tofts et al., 2008). For each b-value, the mixed-signal ADC and corresponding temperature were

$$D_{\mathrm{app}}(b; f) = -\frac{1}{b}\ln\left[\frac{S(b; f)}{S(0)}\right],$$

$$\hat{T}(b; f) = \frac{A}{\ln[B/D_{\mathrm{app}}(b; f)]} - 273.15,$$

with the ACSF calibration $A = 2732$ K and $B = 19.649$ mm$^2$/s (Shibukawa et al., 2021). Because tissue has the smaller diffusion coefficient, its signal decays more slowly and its relative contribution increases with b-value. Tissue admixture therefore lowers the apparent ADC more strongly at higher b-values, producing $\hat{T}(200) > \hat{T}(500) > \hat{T}(700)$ in the CSF-dominated range.

The model observables were the b-dependent endpoint temperature contrast and the downward bias of the primary four-shell median:

$$\Delta T(f) = \hat{T}(200; f) - \hat{T}(700; f),$$

$$B_{\mathrm{PV}}(f) = T_{\mathrm{ref}} - \mathrm{median}_{b\in\{200,500,700,1100\}}\,\hat{T}(b; f), \qquad T_{\mathrm{ref}} = 37.76\ ^\circ\mathrm{C}.$$

$\Delta T(f)$ is zero for both pure CSF and pure tissue. Numerical inversion used the independently selected

CSF-dominated range $0.8 \leq f \leq 1.0$, over which $\Delta T$ increases with tissue signal contribution. Table S2 shows the regenerated lookup values used by the analysis.

Table S2: Regenerated two-compartment lookup table using $D_{\mathrm{CSF}} = 3.0 \times 10^{-3}$ mm$^2$/s, $D_{\mathrm{tissue}} = 0.7 \times 10^{-3}$ mm$^2$/s, and the four-shell temperature estimator. The inferred $f$ is a b = 0 signal fraction, not an anatomical volume fraction.

| CSF b = 0 signal fraction $f$ | $\Delta T(200\text{—}700)$ (°C) | Four-shell median (°C) | Predicted downward bias (°C) |
|---|---|---|---|
| 1.000 | 0.00 | 37.76 | 0.00 |
| 0.995 | 0.16 | 37.46 | 0.30 |
| 0.992 | 0.26 | 37.29 | 0.47 |
| 0.990 | 0.32 | 37.17 | 0.59 |
| 0.980 | 0.63 | 36.60 | 1.16 |
| 0.970 | 0.92 | 36.03 | 1.73 |
| 0.950 | 1.45 | 34.92 | 2.84 |

Within this specified model, $f = 0.95$ produces a 2.84 °C downward bias, comparable in magnitude to the observed 2.36 °C LV–SA difference, but it also predicts $\Delta T(200\text{—}700) = 1.45$ °C. This is 5.9-fold larger than the observed mean SA contrast of 0.245 °C. Conventional tissue admixture large enough to reproduce the regional level difference is therefore incompatible with the weak observed b-dependence under the assumed diffusivities.

For the tissue-admixture sensitivity, the entire positive regional $\Delta T(200\text{—}700)$ was attributed to tissue. At the primary operating point, the group-mean endpoint contrasts were small in both LV and SA (Table S3).

Table S3: Tissue-admixture sensitivity in voxels retained by the primary consistency criterion. $\Delta T$ values are group mean $\pm$ SD across 12 participants; signal fractions and biases are obtained by inverting the group-mean $\Delta T$ through the stated model. Signal fractions are rounded to three decimal places; predicted biases use the unrounded fractions.

| Metric | Lateral ventricles | Subarachnoid space |
|---|---|---|
| $\Delta T(200\text{—}700)$ (°C) | 0.371 $\pm$ 0.224 | 0.245 $\pm$ 0.347 |
| Implied CSF b = 0 signal fraction | 0.988 | 0.992 |
| Model-predicted downward bias (°C) | 0.681 | 0.448 |

The paired LV-minus-SA difference in $\Delta T(200\text{—}700)$ was 0.126 $\pm$ 0.214 °C (95% CI -0.011 to 0.262

°C). Adding the two model-predicted downward biases to their respective regional estimates changed the primary LV–SA difference from 2.357 °C to 2.590 °C. Adding the modelled bias only to SA left 1.909 °C. The biases therefore partly cancelled while preserving the compartment ordering in both modelled sensitivity scenarios. Sensitivity to shell selection, the b = 0 reference, and the noise model is examined in Supplementary Section S3.

Regional measurement-noise uncertainty was propagated through the actual temperature estimator. For each retained voxel $v$, we perturbed its observed four-shell temperatures using the shared-reference model:

$$\tilde{T}_v^{(r)} = \mathrm{median}_i \left( T_{v,i} + \tau_{v,i} Z_0^{(r)} - \eta_{v,i} Z_i^{(r)} \right), \qquad \sigma^2_{\tilde{T},v} = \mathrm{Var}_r \left( \tilde{T}_v^{(r)} \right).$$

We used 16,384 scrambled five-dimensional Sobol draws per voxel, generated using `scipy.stats.qmc.Sobol` (Joe & Kuo, 2008; Matoušek, 1998), with one baseline factor shared across shells and four independent shell factors. The observed temperatures and noise coefficients were plug-in values; the retained population was fixed. The same transformed input-noise reference and actual volume scales used for classification supplied the error coefficients. The regional median-uncertainty scale was

$$\sigma_{R,\mathrm{med}} = \sqrt{\frac{1}{N} \sum_{v=1}^{N} \sigma^2_{\tilde{T},v}}.$$

Using the same approximate nearest-neighbour model, $\rho_{\mathrm{nn}} \approx 0.64$ and three undirected pairs per voxel, the variance-inflation factor was $1 + 2(3)(0.64) = 4.84$. Expressing spatial dependence through an effective sample size follows (Vallejos & Osorio, 2014):

$$N_{\mathrm{eff}} \approx N/4.84, \qquad \mathrm{SE}_{\mathrm{noise,corr}} = \frac{\sigma_{R,\mathrm{med}}}{\sqrt{N_{\mathrm{eff}}}} = \frac{\sqrt{4.84 \sum_v \sigma^2_{\tilde{T},v}}}{N}.$$

The calculation aggregates marginal voxel variances; the nearest-neighbour factor accounts approximately for spatial dependence. The maximum conditional input-reference noise standard errors were 0.044 °C in LV and 0.034 °C in SA.

### S3. Shell and noise sensitivity

The primary voxel temperature was the median across b = 200, 500, 700, and 1100 s/mm$^2$. Because b = 1100 had lower signal-to-noise ratio (SNR) and greater sensitivity to the magnitude-noise floor, we recomputed the regional means using the median of b = 200, 500, and 700 within the same voxel population retained by the primary consistency criterion. Excluding b = 1100 changed the paired difference by -0.05 ± 0.01 °C (Table S4).

Table S4: Sensitivity of regional temperature to excluding b = 1100 from the within-voxel median. Values are group mean ± SD across 12 participants. The consistency classification and retained population remain defined by the primary four-shell analysis.

| Quantity | Four-shell median (°C) | Three-shell median (°C) | Three minus four shells (°C) |
|---|---|---|---|
| Lateral ventricles | 38.41 ± 0.37 | 38.37 ± 0.38 | -0.03 ± 0.01 |
| Subarachnoid space | 36.05 ± 1.10 | 36.07 ± 1.09 | +0.02 ± 0.01 |
| Paired LV–SA difference | 2.36 ± 1.17 | 2.31 ± 1.16 | -0.05 ± 0.01 |

The apparent ADC changed by 1.1% in LV and 0.8% in SA between b = 200 and b = 700 (Table S5). These modest changes indicate residual b-dependence, although the present four-shell acquisition cannot distinguish non-Gaussian diffusion, residual flow, and partial volume.

Table S5: Mean ADC by shell for voxels retained by the primary consistency criterion. ADC is reported as $\times 10^{-3}$ mm$^2$/s; values are group mean ± SD.

| Region | ADC(200) | ADC(500) | ADC(700) | ADC(1100) | $\Delta$ADC$_{200-700}$ |
|---|---|---|---|---|---|
| Lateral ventricles | 3.086 ± 0.048 | 3.050 ± 0.029 | 3.052 ± 0.033 | 3.129 ± 0.043 | 1.1% |
| Subarachnoid space | 2.883 ± 0.094 | 2.863 ± 0.089 | 2.861 ± 0.087 | 2.846 ± 0.107 | 0.8% |

Known-signal simulations assessed the correction after MP-PCA denoising (Table S6). Magnitude data were generated from independent real and imaginary Gaussian noise, using the acquired 109-volume shell counts, then processed with `dwidenoise` (MRtrix3 3.0.8, Exp2, 5 × 5 × 5 patches). We tested CSF temperatures of 36.0 and 38.6 °C, b = 0 amplitude-to-complex-noise ratios of 40–180, and both homogeneous and spatially varying signals, with three independent noise realisations per condition. Temperatures were the four-shell voxel medians, averaged over the 729 interior voxels of each simulated volume.

The quadratic correction produced an upward median-temperature bias; inversion of the magnitude mean and estimated magnitude-noise SD (Koay & Basser, 2006) reduced it. MP-PCA estimates the magnitude mean (Veraart et al., 2016), whereas subtracting $2\sigma^2$ derives from the raw second moment. These operations therefore differ after denoising. The b = 1100 temperature was more sensitive, with quadratic-correction bias ranging from +0.07 to +1.31 °C across individual realisations.

Table S6: Bias of the four-shell median after actual MP-PCA denoising in known-signal Rician simulations. Each interval spans the four condition means (two temperatures, homogeneous or varying signals), each averaged over three noise realisations. Input SNR uses the known complex-channel noise SD.

| Input b = 0 SNR | Quadratic correction bias (°C) | Magnitude-moment inversion bias (°C) |
|---|---|---|
| 40 | +0.165 to +0.198 | +0.016 to +0.029 |
| 60 | +0.087 to +0.103 | -0.002 to +0.006 |
| 80 | +0.055 to +0.063 | -0.003 to -0.001 |
| 120 | +0.029 to +0.033 | -0.002 to +0.002 |
| 180 | +0.015 to +0.018 | -0.001 to +0.002 |

The primary analysis used magnitude-mean inversion with consistent N4 and actual drift scaling of the input-noise reference. A matched quadratic-correction control used the same native MP-PCA result, subsequent processing stages, candidate mask, and anatomical assignments. With consistency selection recomputed separately for each branch, the mean LV–SA difference was 2.44 °C in the quadratic control and 2.36 °C in the primary mean-inversion branch; LV estimates remained higher in every participant. Together with the three-shell summary, these comparisons support robustness of the regional ordering. The Rician simulations in Table S6 evaluate the denoising stage; acquisition metadata supported the Rician reconstruction assumption.

#### b = 0 reference sensitivity

The four primary shell-specific temperature estimates used the same average of 11 b = 0 volumes. This creates a shared random contribution across shells, which is represented explicitly in the noise-only consistency null. We tested whether regional temperatures or the paired LV–SA difference depended materially on the choice of b = 0 reference. All alternatives were evaluated in the retained LV and

SA voxel populations from the primary analysis; clustering labels, anatomical adjudication, and voxel inclusion were held fixed so that the comparison isolated the temperature estimator.

Four alternatives were compared with the scan-wide mean b = 0 reference. First, a time-local reference was linearly interpolated from the b = 0 acquisitions bracketing each diffusion-weighted acquisition and then averaged within shell. Second, the 11 b = 0 volumes were split among the four shells in balanced groups of 3, 3, 3, and 2 across 512 deterministic allocations; this deliberately inefficient analysis was an independence stress test. Third, ADC was estimated directly from the b = 200 and 500 signals. Fourth, a free-intercept log-linear fit

$$\ln S(b) = \alpha - bD$$

was applied to b = 200, 500, and 700, removing explicit dependence on b = 0 and on any multiplicative signal scale common to those shells. Every estimator had 100% valid coverage of the fixed voxel populations.

Table S7: Sensitivity of regional temperatures to the b = 0 reference. Values are group mean ± SD across 12 participants. All estimates use the fixed retained voxel populations from the primary analysis; the LV–SA difference was positive in all 12 participants for every estimator.

| Estimator | LV temperature (°C) | SA temperature (°C) | LV–SA difference (°C) |
|---|---|---|---|
| Global b = 0 | 38.41 ± 0.37 | 36.05 ± 1.10 | 2.36 ± 1.17 |
| Time-local b = 0 | 38.42 ± 0.39 | 35.95 ± 1.05 | 2.47 ± 1.11 |
| Disjoint b = 0 | 38.49 ± 0.37 | 36.01 ± 1.15 | 2.48 ± 1.19 |
| b = 200–500 | 38.03 ± 0.28 | 35.90 ± 1.14 | 2.13 ± 1.14 |
| Free-intercept b = 200/500/700 | 38.17 ± 0.32 | 35.92 ± 1.08 | 2.25 ± 1.15 |

Relative to the primary estimator, the mean changes in the LV–SA difference were +0.11 °C for the time-local reference, +0.12 °C for the disjoint reference, -0.23 °C for b = 200–500, and -0.11 °C for the free-intercept b = 200/500/700 fit. Across the 512 balanced disjoint allocations, the central 95% of group-mean differences was approximately 2.39–2.56 °C. For that estimator, each participant's regional values and paired difference were summarised separately by their allocation medians; the paired value need not equal the difference of the two regional medians. The free-intercept analysis provided the strongest test of explicit b = 0 dependence: its 2.25 ± 1.15 °C difference was positive in every participant and close to the primary 2.36 ± 1.17 °C estimate.

The temporal coefficient of variation of b = 0 signal averaged 1.33% in LV and 2.34% in SA. Mean fitted end-to-end signal change was -0.35% in LV and -3.34% in SA. This spatially different change could reflect residual motion, changing partial-volume composition, or other acquisition effects rather than scanner drift. Time-local referencing changed LV by +0.02 °C, SA by -0.10 °C, and the paired difference by +0.11 °C.

These comparisons assessed temperature estimators within the primary selected population. The scan-wide average of all 11 b = 0 volumes retains the full reference precision, with its shared noise contribution represented in the consistency null.

#### Residual-bootstrap robustness

As a model-assisted sensitivity analysis, we resampled the centred, finite-sample-corrected residuals of the acquired volumes within each voxel. Each bootstrap replicate used one resampled b = 0 mean shared across all four shell-temperature estimates and independent residual resamples within the four diffusion shells. The null statistic was the cross-shell MAD of the bootstrap temperature errors $(T^* - T)$. All eligible voxels received 2,000 replicates; voxels near a classification boundary were adaptively refined with 10,000–50,000 replicates.

For the direction-adjusted variant, a fitted trace-free $l = 2$ directional component was removed within each shell before residual resampling, with a correction for the regression residual degrees of freedom. The primary analytic model remained the principal measurement-noise model; the two bootstrap variants were broader robustness checks (Table S8).

The analytic model propagates the N4- and drift-scaled MP-PCA input-noise reference. The residual-bootstrap alternatives also capture across-volume physiological and preprocessing variation.

Table S8: Sensitivity of retained-voxel count and regional temperature to residual-based null models. Retained-voxel counts are pooled across participants. Temperatures and differences are group mean ± SD across 12 participants; confidence intervals are for the paired LV–SA difference.

| Model | Pooled retained LV voxels | Pooled retained SA voxels | LV temperature (°C) | SA temperature (°C) | LV–SA difference (°C; 95% CI) | Paired $p$ |
|---|---|---|---|---|---|---|
| Primary analytic | 22,476 | 18,367 | 38.407 ± 0.372 | 36.050 ± 1.103 | 2.357 ± 1.169 (1.614–3.100) | $2.32 \times 10^{-5}$ |

| Model | Pooled retained LV voxels | Pooled retained SA voxels | LV temperature (°C) | SA temperature (°C) | LV–SA difference (°C; 95% CI) | Paired $p$ |
|---|---|---|---|---|---|---|
| Direction-adjusted residual bootstrap | 21,750 | 34,505 | 38.623 $\pm$ 0.393 | 35.885 $\pm$ 1.080 | 2.737 $\pm$ 1.144 (2.011–3.464) | $4.65 \times 10^{-6}$ |
| Raw residual bootstrap | 25,177 | 36,733 | 38.573 $\pm$ 0.395 | 35.866 $\pm$ 1.096 | 2.707 $\pm$ 1.158 (1.971–3.442) | $5.81 \times 10^{-6}$ |

For the primary analytic model, Cohen's $d_z$ for the paired LV–SA difference was 2.02. Both bootstrap variants preserved higher LV than SA estimates in all 12 participants and retained substantially more SA voxels than the analytic model. Their selected populations differed from the primary population; the direction-adjusted variant retained slightly fewer LV voxels. Across-volume residuals can contain directional signal, physiology, motion, interpolation, and preprocessing variation in addition to measurement noise. These analyses support robustness of the regional ordering while showing sensitivity of mask size and the difference magnitude to the assumed null.

After final 50,000-replicate refinement, 100 raw-bootstrap and 103 direction-adjusted voxel classifications differed between independent seeds among 350,333 eligible LV and SA voxels (four-shell valid with median temperature at least 25 °C). Allowing all resolutions of these boundary cases gave group-difference envelopes of 2.704–2.708 °C and 2.735–2.740 °C, respectively, and preserved every participant's LV–SA ordering. These envelopes quantify finite-simulation ambiguity.

### S4. SA compartment size and tympanic temperature

SA temperature increased with the number of voxels retained by the primary consistency criterion (Spearman $\rho = +0.76$; fixed-seed Monte Carlo permutation $p = 0.006251$ from 999,999 permutations). The absolute SA–ear discrepancy decreased as retained SA voxel count increased ($\rho = -0.66$; permutation $p = 0.023718$). Restricting the cohort to progressively larger retained compartments increased the SA mean and reduced its between-participant standard deviation, whereas the post-session tympanic mean remained stable (Supplementary Figure S1 and Table S9). In the six largest compartments, mean SA temperature was 36.86 °C compared with 36.80 °C for tympanic temperature, giving a paired SA–ear difference of $+0.06 \pm 0.49$ °C (95% CI -0.45 to $+0.57$ °C; paired t-test $p = 0.778$). Together with the continuous count–temperature association, these post hoc comparisons support prospective evaluation of regional sampling criteria based on retained count, effective sample size, and regional uncertainty.

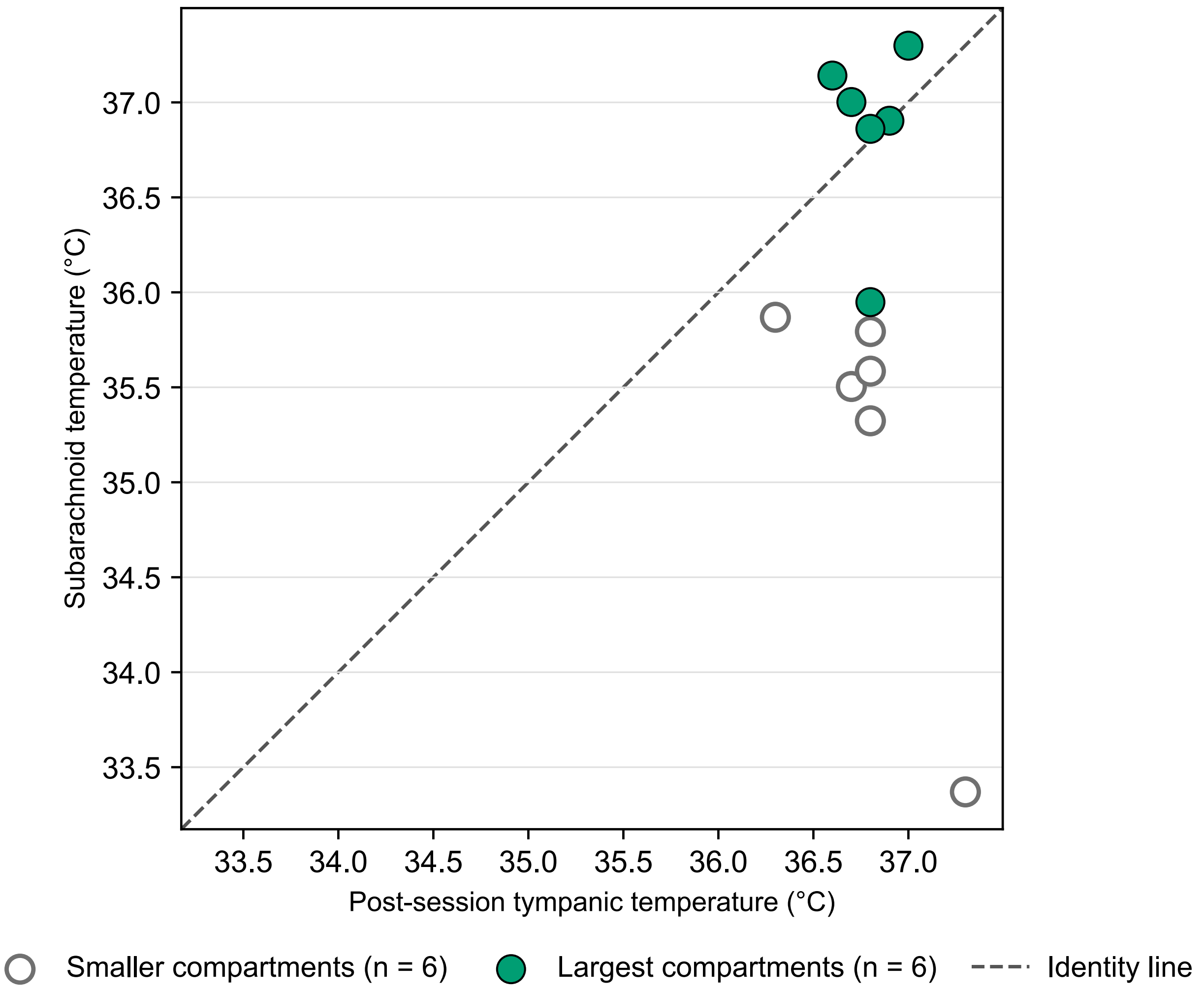


**Supplementary Figure S1. Subarachnoid temperature is closer to post-session tympanic temperature in the largest retained compartments.** Participant-level subarachnoid temperature plotted against post-session tympanic temperature. The dashed diagonal is the identity line; the post hoc subset of the six participants with the largest retained subarachnoid voxel counts is highlighted in green. Their mean subarachnoid–ear difference was $+0.06 \pm 0.49$ °C. Together with the continuous association between subarachnoid temperature and retained voxel count in main Figure 4c, this convergence motivates prospective evaluation of sampling-based quality criteria for regional subarachnoid thermometry.

Table S9: SA and post-session tympanic temperature in nested post hoc cohorts defined by retained SA voxel count.

| Subset | n | Minimum retained SA voxels | $T_{\mathrm{SA}}$ (°C) | Post-session $T_{\mathrm{ear}}$ (°C) | $T_{\mathrm{SA}} - T_{\mathrm{ear}}$ (°C) |
|---|---|---|---|---|---|
| All | 12 | 295 | 36.05 $\pm$ 1.10 | 36.79 $\pm$ 0.24 | -0.74 $\pm$ 1.22 |
| Top 8 | 8 | 960 | 36.57 $\pm$ 0.68 | 36.73 $\pm$ 0.21 | -0.16 $\pm$ 0.61 |
| Top 6 | 6 | 1,162 | 36.86 $\pm$ 0.47 | 36.80 $\pm$ 0.14 | +0.06 $\pm$ 0.49 |

### S5. Comparison with FA filtering

We compared the post-processing criterion with the fractional anisotropy (FA) threshold used by Shibukawa et al. (2021). FA was refitted from the revised b = 0, 200, 500, 700, and 1100 data using MRtrix3 weighted least squares with two iterative reweighting steps; b = 1500 was excluded. FA $<$ 0.1 was applied within the fixed anatomical compartments, and temperature was estimated at b = 500 s/mm$^2$. Classification overlap was calculated among anatomically assigned candidates with finite FA; every primary retained voxel was represented. This comparison evaluates the two post-processing criteria within our conventional bipolar acquisition; Shibukawa et al. used second-order motion-compensated encoding.

The criteria test different signal properties. Low FA indicates that the measured diffusion is approximately isotropic, but it does not establish that a voxel contains only freely diffusing CSF. In particular, admixture with approximately isotropic tissue can remain below an FA threshold while changing the apparent diffusion coefficient across b-values. The multi-shell criterion instead tests whether the four temperature estimates agree within the dispersion expected from voxel-specific measurement noise.

Low FA was not sufficient to identify cross-shell-consistent measurements (Supplementary Figure S2). Among voxels passing FA $<$ 0.1, 70.8% in LV and 92.2% in SA failed the multi-shell consistency criterion. Conversely, 85.0% of consistency-passing LV voxels and 93.8% of consistency-passing SA voxels also passed FA $<$ 0.1. Among finite-FA anatomical candidates, 17.5% of LV voxels passed both criteria, 42.4% passed FA alone, 3.1% passed consistency alone, and 37.0% passed neither. The corresponding SA fractions were 6.9%, 81.9%, 0.5%, and 10.7%. The FA-filtered b = 500 estimate was lower than the primary multi-shell estimate by a mean paired difference of 0.76 °C in LV (37.65 versus 38.41 °C) and 1.96 °C in SA (34.09 versus 36.05 °C). This colder shift is compatible with the inclusion of tissue-contaminated measurements, particularly in SA. Cross-shell disagreement can also reflect residual motion, flow, or other violations of the free-diffusion model.

Second-order motion compensation and multi-shell consistency therefore address complementary limitations. Motion-compensated encoding reduces acquisition sensitivity to coherent CSF motion, including motion effects that a cross-shell agreement test cannot detect if they shift all shell estimates similarly, but it does not address tissue partial volume. The multi-shell criterion detects b-dependent deviations that may remain approximately isotropic and pass FA filtering. Combining motion-compensated encoding with multi-shell consistency testing would therefore address complementary acquisition- and post-processing-level sources of error.

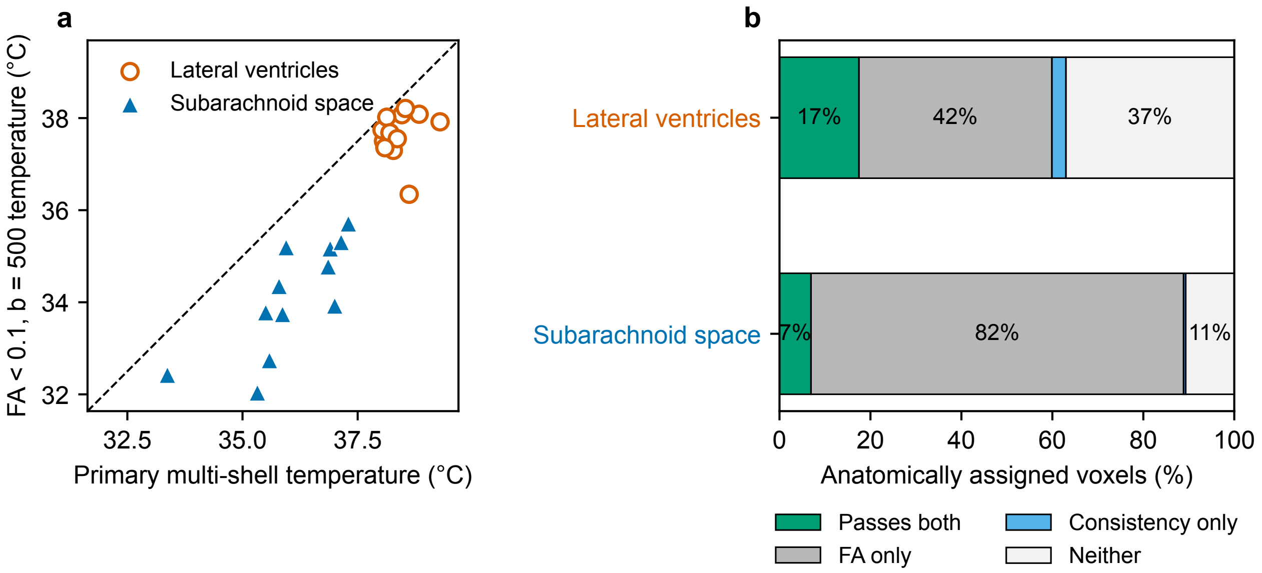


**Supplementary Figure S2. Low FA does not ensure cross-shell-consistent CSF thermometry.** Panel a shows participant-level temperature obtained with FA $< 0.1$ and single-shell b $=$ 500 s/mm$^2$ thermometry, plotted against the primary multi-shell estimate. Open orange circles show the lateral ventricles and filled blue triangles show the subarachnoid space; the dashed line is identity. Panel b shows voxel-classification overlap among finite-FA candidates within the anatomical compartments, pooled across participants. Of the voxels passing FA $< 0.1$, 70.8% in the lateral ventricles and 92.2% in the subarachnoid space failed the multi-shell consistency criterion. The comparison evaluates the FA threshold on our bipolar acquisition.

### S6. Synthetic digital reference demonstration

We created a deterministic BIDS-compatible digital reference dataset to test the public implementation against known inputs. The dataset contains no participant-derived images, metadata, anatomical template, or fitted participant parameters. Analytic geometry defines lateral-ventricular (LV), subarachnoid (SA), and artefact-rich Other compartments. The acquisition contains 109 volumes: 11 b = 0 volumes, 20 directions at b = 200, 500, and 700 s/mm$^2$, and 19 directions at b = 1100 and 1500 s/mm$^2$. As in the primary analysis, the validation used b = 200, 500, 700, and 1100 s/mm$^2$; b = 1500 s/mm$^2$ was present but excluded.

All 109 magnitude volumes underwent MP-PCA denoising (Exp2, 5 $\times$ 5 $\times$ 5 patches), followed by Rician magnitude-moment inversion before shell averaging, including b = 0. The estimated input magnitude-noise SD was used for propagation. The bias-field and drift scale factors were unity. Classification used the primary voxel-specific shared-b = 0 null and adaptive draw protocol. LV and SA were the primary eligible compartments; Other was evaluated separately with the same rule as a shell-inconsistency challenge and did not enter regional means. HDBSCAN described geometry and truth overlap.

The programmed LV and SA temperatures were 38.8 °C and 36.3 °C, respectively, giving an LV–SA difference of 2.50 °C. Other had shell-dependent apparent temperatures of 34–52 °C. The primary example recovered retained-voxel means of 38.80 °C in LV and 36.35 °C in SA, giving a difference of 2.45 °C. It retained 741/744 LV voxels and 3,068/7,698 SA voxels; none of the 816 Other challenge voxels passed. HDBSCAN returned four truth-pure clusters, with the two LV halves represented separately.

A matched quadratic-correction control used the same denoised data and estimated input-noise reference, with selection recomputed. It recovered an LV–SA difference of 2.46 °C and retained 741 LV and 3,736 SA voxels; no Other challenge voxels passed. These known-input results verify the released calculation and recovery of the programmed temperatures under the specified simulation conditions.

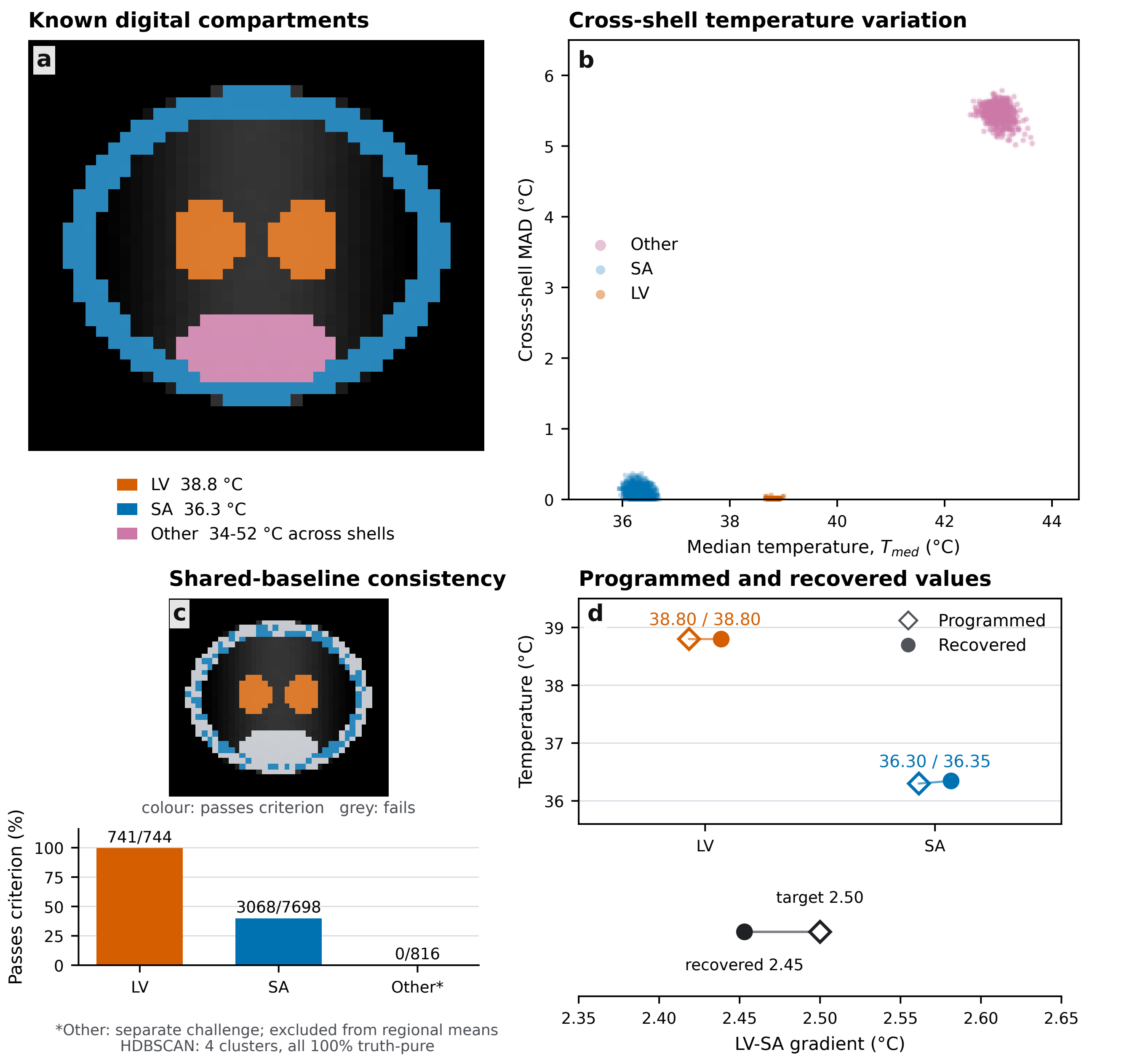


**Supplementary Figure S3. Recovery of programmed temperatures and cross-shell consistency in an analytic synthetic dataset. a**, Known digital CSF compartments overlaid on the corrected synthetic b = 0 image. LV and SA have fixed programmed temperatures; Other has shell-dependent apparent temperatures. **b**, Saved corrected voxel-wise median temperature, $T_{\mathrm{med}} = \tilde{T}$, and cross-shell MAD from b = 200, 500, 700, and 1100 s/mm$^2$. **c**, Spatial decisions and compartment-specific fractions passing the voxel-specific shared-baseline median-null criterion; colour denotes passing voxels and grey denotes failing voxels. Other is a separate same-rule challenge and is excluded from regional estimates. HDBSCAN diagnoses geometry/truth overlap. **d**, Programmed and recovered retained-voxel LV/SA means and their difference. No participant-derived data are used.
